# Plasticity in striatal dopamine release is governed by release-independent depression and the dopamine transporter

**DOI:** 10.1101/392753

**Authors:** Mark D. Condon, Nicola J. Platt, Yan-Feng Zhang, Bradley M. Roberts, Michael A. Clements, Stefania Vietti-Michelina, Min-Yee Tseu, Katherine R. Brimblecombe, Sarah Threlfell, Ed O. Mann, Stephanie J. Cragg

## Abstract

Mesostriatal DA neurons possess extensively branched axonal arbours. Whether action potentials are converted to DA output in striatum will be influenced dynamically and critically by axonal properties and mechanisms that are poorly understood. We addressed the roles for mechanisms governing release probability and axonal activity in determining short-term plasticity of DA release, using fast-scan cyclic voltammetry in *ex vivo* mouse striatum. Brief short-term facilitation (STF) and longer short-term depression (STD) were only weakly dependent on the level of initial release, i.e. were release-insensitive. Rather, short-term plasticity was strongly determined by mechanisms which governed axonal activation, including K^+^-gated excitability and the dopamine transporter (DAT), particularly in dorsal striatum. We identify the DAT as a master regulator of DA short-term plasticity, governing the balance between release-dependent and independent mechanisms that also show region-specific gating.

**Key Findings:**

- Short-term plasticity in dopamine release is only weakly governed by initial release
- Short-term depression is strongly dependent on axonal excitability and activation
- The dopamine transporter controls short-term plasticity and drives short-term depression
- Dopamine transporters govern the balance between release-dependent and -independent mechanisms

## Introduction

Dopamine (DA) release in the striatum plays key roles in action selection and behavioural reinforcement, and is dysregulated in diverse disorders including Parkinson’s disease and addictions. DA neurons generate action potentials at a range of frequencies, firing tonically below 10 Hz, and intermittently in short bursts with instantaneous frequencies of up to approximately 40 Hz [1-6]. Bursts occur in response to salient environmental stimuli predicting reward value or action signals [6-8]. However, the extent to which action potential activity is conveyed into striatal DA release remains unclear; DA axons are important sites for local regulation of DA output [9-13] which shows strong short-term plasticity and distorts the relationship between activity and DA output [11,14-17].

Despite classical accounts of axons as simple high-fidelity cables, neurotransmitter release following action potentials in CNS neurons is shaped by factors that regulate action potential propagation and axonal excitability such as axonal morphology and branching, the expression of ion channels and presynaptic receptors, and other diverse mechanisms that regulate vesicle pools, neurotransmitter release probability and short-term plasticity [18]. Axons of DA neurons are remarkable: they comprise vast, extensively branched arbours [19, 20], that from anatomical observations and a binary tree model [21] can be calculated to form an average of 16,000 branch points per nigrostriatal neuron. Axonal properties are likely to be particularly important in governing axonal activity and striatal DA output. One major influence is the input from striatal cholinergic interneurons onto nicotinic receptors (nAChRs) on DA axons, which influence the pattern of DA release and promote short-term depression (STD) [11,15,22-24]. But even in the absence of nAChR activation, DA release shows an underlying spectrum of short-term plasticity ranging from short-term depression (STD) to short-term facilitation (STF) [11,15,23,25,26] for which the underlying mechanisms are poorly understood. Existing evidence suggests that short-term plasticity might be only weakly related to initial levels of release [17,25,26] but the dominant drivers remain undefined.

Here, we delineate the roles of three types of intrinsic drivers that could underlie short-term plasticity of DA release from striatal DA axons. We examine, firstly, the role for initial release; secondly, the role for K^+^-dependent mechanisms that will govern overall excitability and repolarization; and thirdly, a potential role for the dopamine transporter (DAT). Besides mediating DA uptake, the DAT generates a depolarising transport-coupled conductance in midbrain DA neurons [27, 28], and has been suggested to limit vesicular release [29-32]. Thus, the DAT has the potential to govern short-term plasticity of DA release via regulation of axonal activation and/or release probability (P_r_).

We reveal that initial release plays a limited role in short-term plasticity except on the shortest of timescales (10 - 25 ms), when STF operates. The prevailing STD at longer timescales (25 ms – 200 ms) is apparently independent of the level of initial release i.e. it is “release-insensitive”. We identify that STD is strongly determined by mechanisms that influence membrane activation, particularly in dorsal striatum. Furthermore, we identify that the DAT drives STF at shortest intervals and STD at longer, physiological inter-pulse intervals, effectively clamping release. We propose a region-specific hierarchy of interacting drivers of short-term plasticity, overseen by the DAT, with the DAT promoting release-insensitive over release-dependent mechanisms.

## Results

### Patterns of short-term plasticity at DA release sites are region-specific and weakly release-dependent

We assessed short-term plasticity of DA release in acute coronal striatal slices evoked using pairs of local electrical pulses and inter-pulse intervals (IPI) of 10-200 ms in both dorsolateral striatum (caudate-putamen, CPu) and nucleus accumbens core (NAc). Confounding effects of nAChRs were excluded by inclusion throughout of antagonist DHβE (see Methods). The ratio of DA release evoked by the second pulse in a pair compared to a single pulse (paired-pulse ratio, PPR) varied with IPI in CPu (Fig. 1A,C) and in NAc (Fig. 1B,C). In both regions, PPR decreased with increasing IPI. Short-term facilitation (STF, PPR>1) was observed at the shortest IPIs (10 ms in CPu, 10-25 ms in NAc), whereas strong short-term depression (STD) (PPR<1) was observed at longer IPIs in both regions (≥ 25 ms in CPu, >40 ms in NAc), as shown previously [15, 33]. In CPu, STD was significantly more pronounced than in NAc (Fig. 1C, Two-way ANOVA, region x IPI interaction, F_4,25_ = 4.195, *P* = 0.010, *n* = 7) suggesting region-specific gating of short-term plasticity.

PPR at classic fast synapses is typically inversely related to initial P_r_, resulting in STF when P_r_ is low, and STD when P_r_ is high [34-37]. STF at these synapses consists of several temporally-overlapping processes with a range of time constants from 30 – 500 ms [38]. An inverse relationship between PPR and single pulse release has been reported in CPu for short IPI of 10 ms but not 100 ms [25], suggesting that short-term plasticity of DA release might reflect initial P_r_ at only very short intervals. We probed the relationship between PPR and the extracellular concentration of DA ([DA]_o_) evoked by a single pulse (as a proxy of initial P_r_) in CPu and in NAc across a population of release sites that exhibited a 10-fold range in [DA]_o_ evoked by a single pulse (1p [DA]_o_). We found an inverse relationship between 1p [DA]_o_ and PPR for IPI of 10 ms in both CPu and NAc, and at 40 ms in NAc only, but in neither region at the longer interval of 100 ms (Fig. 1D, linear regressions, 10 ms, CPu: β = -0.79 ± 0.14 [95% CI: -1.08 to -0.50], F_1,33_ = 30.9, *P* = 3.56 × 10^-6^, *n* = 35; NAc: β = - 0.52 ± 0.19 [95% CI: -0.93 to -0.11], F_1,17_ = 7.136, *P* = 0.016, *n* = 19; 40 ms, CPu, *P* = 0.152, *n* = 35; NAc 95% CI: -0.38 to -0.001], F_1, 17_ = 4.489, *P* = 0.049, *n* = 19; 100 ms, CPu: *P* = 0.122, n = 35; NAc: *P* = 0.917, n = 19). These data indicate that DA PPR is a function of P_r_ at only very short intervals and specifically, only at those intervals at which STF can be observed. These data suggest that short-term plasticity in DA release might be governed by release-dependent mechanisms at very short intervals, but by mechanisms unrelated to initial release at longer more physiological intervals.

**Figure 1.**
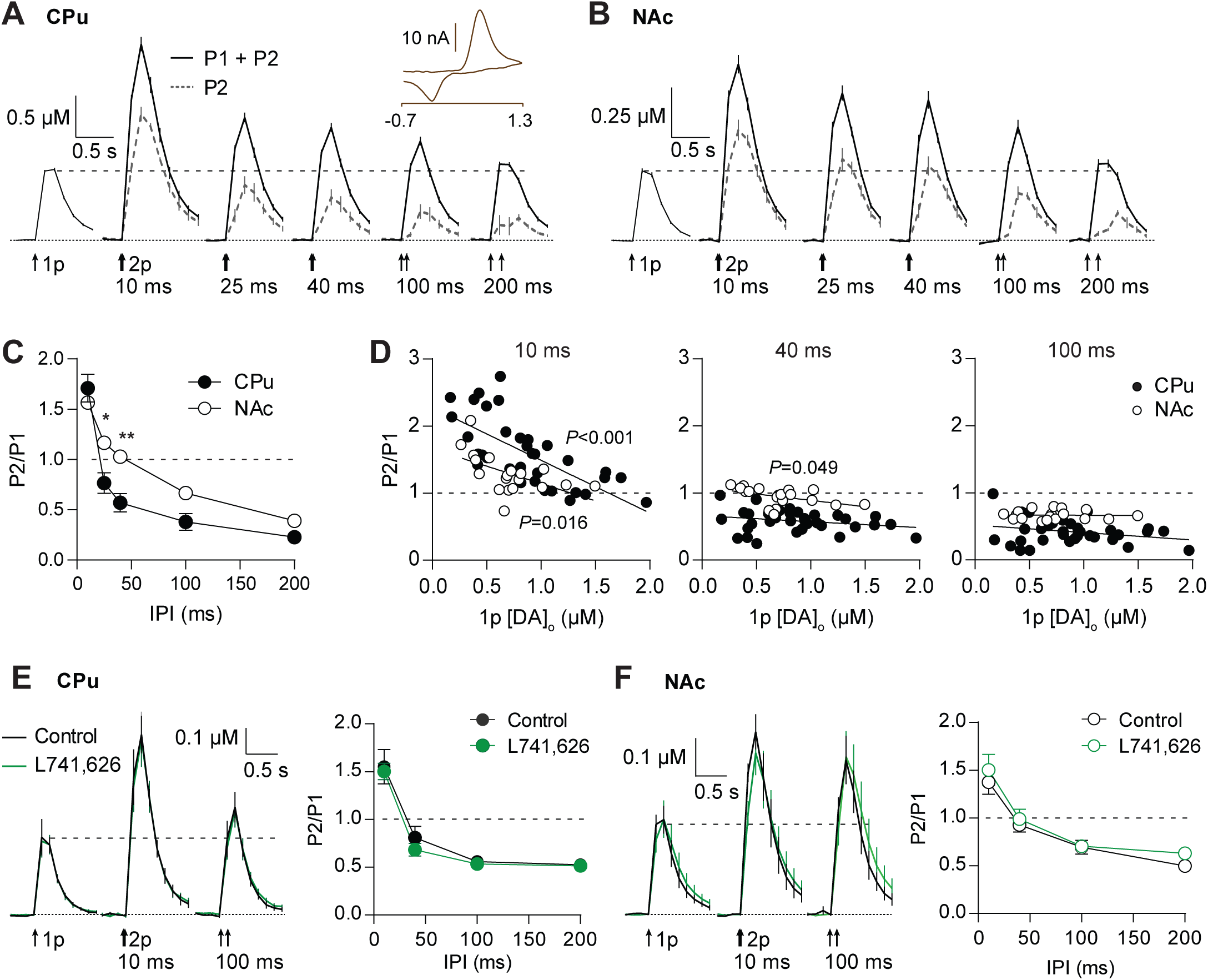
Short-term plasticity at DA release sites is region-specific and only weakly release-dependent. **(A, B)** Mean profiles of [DA]_o_ transients (± SEM) elicited by single or paired electrical pulses (*arrows*) at IPI of 10-200 ms in CPu (A, *n* = 4) and NAc (B, *n* = 3). Dotted traces, [DA]_o_ attributable to the second pulse, P2, in a pair (P1+P2 – P1). *Inset*, representative voltammogram for evoked DA release. **(C)** Mean P2/P1 (± SEM) vs. IPIs in CPu (*filled circles*) and NAc (*open circles*). Where error bars are not visible they fall within the bounds of the symbol. **(D)** P2/P1 vs peak 1p-evoked [DA]_o_ for IPIs of 10 ms (*left*), 40 ms (*centre*) and 100 ms (*right*). Each data point is 1 site in CPu (*filled circles*, *n* = 35) or NAc (*open circles*, *n* = 19). Linear fits to data show *P*-values (F-test) to indicate whether the slope is significantly > 0. **(E,F)** (*Left)* Mean profiles of [DA]_o_ transients in CPu (E, *n* = 5) and NAc (F, *n* = 6) in control conditions vs L741,626 (1 µM, *green*). (*Right*) Mean P2/P1 vs IPIs in control conditions and L741,626 in CPu (E) or NAc (F). Two-way ANOVA with Bonferroni’s test for *post-hoc* comparisons: \**P* < 0.05, \*\**P* < 0.01. Source data are provided as a Source Data file.

This expression of STD was not due to activation of inhibitory D_2_-autoreceptors (D2Rs). Antagonism of D2Rs with antagonist L741,626 (1 µM), did not influence peak [DA]_o_ (Fig. 1E) or PPR at IPIs up to 200 ms in either CPu or NAc (Fig. 1F, Two-way ANOVA, CPu: *P* = 0.420, n= 5, NAc: *P* = 0.2197, n = 6), although did for longer IPIs (**Supp. Fig S1**) consistent with previous findings that D2Rs only modify release after longer IPIs (>200 ms, < 3 s) and are not tonically regulating DA release in brain slices [14,39-41]. We also ruled out an alternative effect whereby DA remaining after the first pulse might determine the release by the second, with higher DA remaining at short IPI driving STF and lower DA at longer IPI driving STD. We varied the intensity of the initial electrical stimulus to halve the [DA]_o_ evoked in CPu, but showed that the [DA]_o_ evoked by a second pulse at a fixed IPI of 25 ms and a fixed stimulus strength released the same [DA]_o_ at P2 regardless of initial level of [DA]_o_ (**Suppl. Fig S2**).

### Short-term plasticity is shaped by calcium-dependent facilitation and release-insensitive depression

To test directly whether short-term plasticity can be shaped by changes to P_r_, we varied initial release at a given release site by varying the extracellular concentration of Ca^2+^ ([Ca^2+^]_o_) in a 3-fold range (1.2 - 3.6 mM). [DA]_o_ evoked by 1p was strongly dependent on [Ca^2+^]_o_, in both CPu and NAc (Fig. 2A,B) as expected. There was a steeper relationship between [Ca^2+^]_o_ and 1p [DA]_o_ in CPu than in NAc (Fig. 2C, Two-way ANOVA, [Ca^2+^]_o_ x region interaction, F_2,20_ = 17.25, *P* = 4.43 x10^-5^, *n* = 6), consistent with previous findings [26]. However, despite large changes in 1p-evoked DA release in both regions (4-fold in CPu, 2.5-fold in NAc), [Ca^2+^]_o_ had only modest effects on PPR limited to very short intervals. In CPu, a trend towards an inverse relationship at 10 ms IPI did not reach significance (Fig. 2D,F, Two-way ANOVA, *P* = 0.604, n = 4). In NAc, PPR varied inversely with [Ca^2+^]_o_ but at only the shortest interval of 10 ms IPI (Fig. 2E,F, Two-way ANOVA, IPI x [Ca^2+^]_o_ interaction, F_8,30_ = 4.497, *P* = 0.001, *n* = 3). Furthermore, we noted that the size of effect on PPR in NAc versus CPu did not tally with the size of effect of [Ca^2+^]_o_ on 1p [DA]_o_: the range of 1p [DA]_o_ were less in NAc than CPu despite the larger range in short-term plasticity (Fig. 2F). The difference in short-term plasticity between CPu and NAc cannot therefore be attributed to a difference in the sensitivity of initial P_r_ to [Ca^2+^]_o_ (see Fig. 2C). Together these data show that short-term plasticity of DA release is not easily explained by Ca^2+^-limited mechanisms, suggesting that other mechanisms dominate that are independent of initial release.

**Figure 2.**
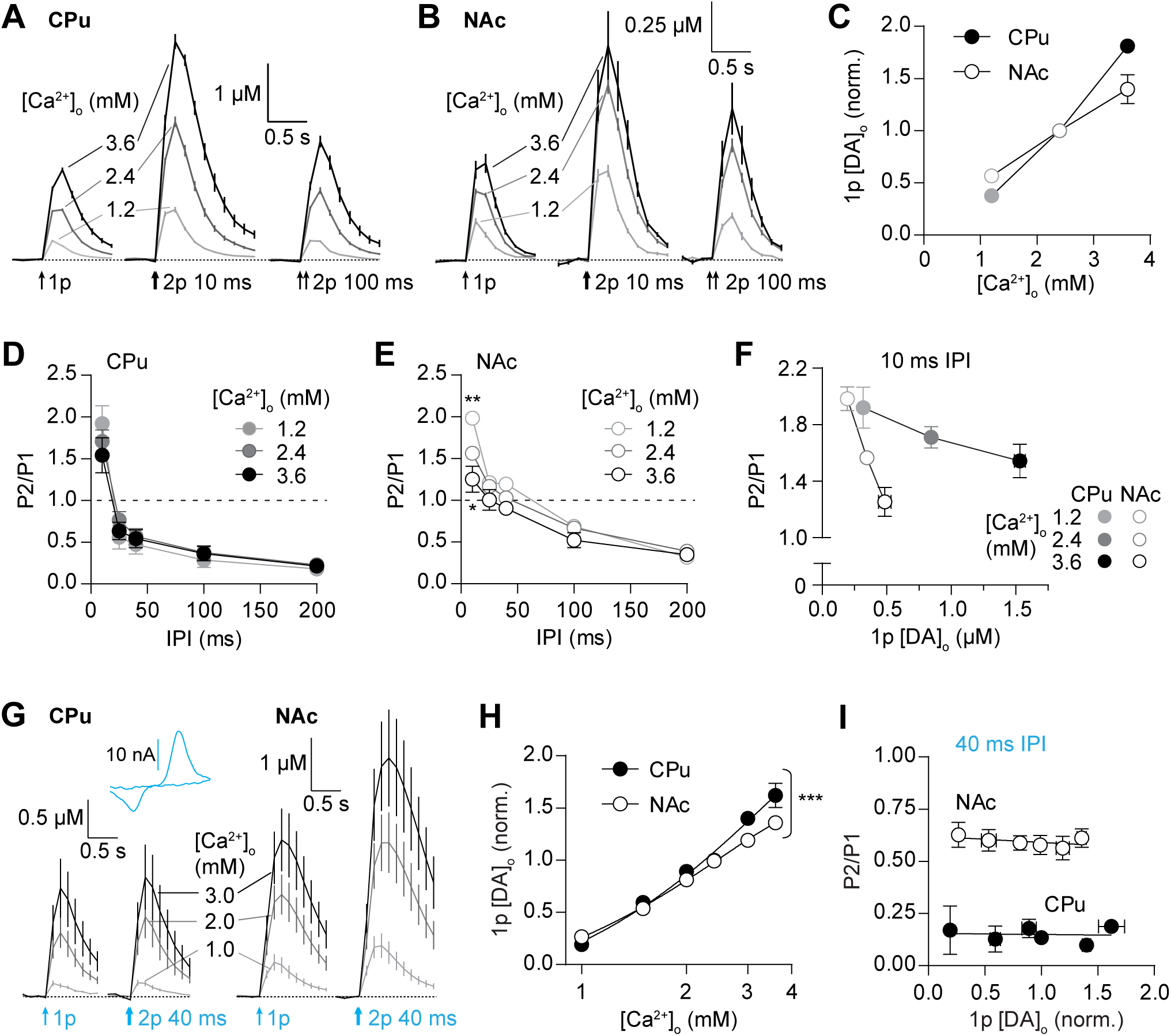
Short-term plasticity is shaped by calcium-dependent facilitation and release-independent depression. **(A, B)** Mean profiles of [DA]_o_ transients (± SEM) evoked by single or paired electrical pulses in 1.2 mM (*light grey*), 2.4 mM (*dark grey*) and 3.6 mM (*black*) [Ca^2+^]_o_ in CPu (A, *n* = 3) and NAc (B, *n* = 3). **(C)** Mean 1p evoked [DA]_o_ (± SEM, normalised to 2.4 mM [Ca^2+^]_o_) vs [Ca^2+^]_o_ in CPu (*filled*) and NAc (*open circles*). **(D, E)** Mean P2/P1 (± SEM) vs. IPI in 1.2 mM (*light grey*), 2.4 mM (*dark grey*) and 3.6 mM (*black*) [Ca^2+^]_o_ in CPu (D, *filled*) and NAc (D, *open circles*). **(F)** Mean P2/P1 (± SEM) vs mean peak 1p [DA]_o_ (± SEM) at 10 ms IPI in 1.2 mM (*light grey*), 2.4 mM (*dark grey*) and 3.6 mM (*black*) [Ca^2+^]_o_ in CPu (*filled*) and NAc (*open circles*). **(G)** Mean profiles of [DA]_o_ transients (± SEM) elicited by single or paired light pulses at 40 ms IPI (473 nm, 2 ms) in 1.0 mM (*light grey*), 2.0 mM (*dark grey*) and 3.0 mM (*black*) [Ca^2+^]_o_ in CPu (*left*, *n* = 3) and NAc (*right*, *n* = 3). *Inset*, representative voltammogram for optogenetically voked DA release. **(H)** Mean peak 1p [DA]_o_ (± SEM, normalised to 2.4 mM [Ca^2+^]_o_) vs [Ca^2+^]_o_ (log scale) evoked by light stimulation in CPu (*filled*) and NAc (*open circles*). Second-order polynomial curve fit; F-test of relative sum of squares shows curve fits are different. **(I)** Mean P2/P1 (± SEM) at 40 ms IPI vs mean peak 1p [DA]_o_ (± SEM) evoked by optical stimulation in CPu (*filled*) and NAc (*open circles*). Two-way ANOVA with Bonferroni’s test for *post-hoc* comparisons except where stated otherwise: \**P* < 0.05, \*\**P* < 0.01, \*\*\**P* < 0.001. Source data are provided as a Source Data file.

To confirm that STD and its insensitivity to [Ca^2+^] do not result from stimulation of an undefined input to DA axons, we used targeted optogenetic stimulation of DA axons. In striatum from heterozygote DAT^IRES-Cre^ mice expressing ChR2-eYFP after viral delivery, a brief light flash (2 ms, 473 nm wavelength) evoked [DA]_o_ transients that varied with [Ca^2+^]_o_ (Fig. 2G) and with a steeper [Ca^2+^]_o_ concentration-response curve in CPu than in NAc (Fig. 2H, nonlinear regression, CPu: R^2^ = 0.91, n = 3, NAc: R^2^ = 0.94, n = 3, comparison of fits: F_4,118_ = 5.94, *P* = 2 × 10^-4^), as seen for electrically evoked DA release. Furthermore, at an interval of 40 ms, at which ChR2 can reliably drive repeat spiking in DA neurons [1] and DA release [11], PPR was greater in NAc than in CPu, as observed using electrical stimulation (see Fig. 1C). However, even with optogenetic stimulation, there was no inverse relationship between PPR and 1p [DA]_o_ (Fig. 2I, linear regression, CPu: *P* = 0.874, n = 3, NAc: *P* = 0.286, n = 3). These data confirm that STD is not attributable to stimulation of other inputs. For subsequent experiments, we chose electrical over optogenetic stimulation to avoid the potentially confounding effects of Ca^2+^ entry through ChR2 on P_r_ and short-term plasticity.

### Release-independent STD is controlled by K^+^-dependent gating

We tested whether STD was gated by axonal membrane excitability by varying [K^+^]_o_. Varying [K^+^]_o_ can alter membrane potential and repolarisation through the Nernstian driving forces underlying K^+^-mediated currents [42] and also through K^+^-dependent inhibition of K^+^-channel inactivation [43]. Reduced [K^+^]_o_ can thereby favour membrane hyperpolarisation and faster repolarisation between pulses, promoting de-inactivation of Na^+^ channels, but can also promote inactivation of K^+^-channels, leading to use-dependent depolarisation and enhanced Na^+^-channel recruitment. In either case, recruitment of voltage-gated Na^+^ channels at a subsequent depolarising event will be altered.

We found that varying [K^+^]_o_ (1.25 - 7.5 mM) did not change 1p [DA]_o_ in either CPu (Fig. 3A) or NAc (Fig. 3B) but nonetheless significantly modulated short-term plasticity (Fig. 3A-D). In CPu, PPR varied inversely with [K^+^]_o_ across IPIs, with STD minimized at lowest [K^+^]_o_ (Fig. 3C, Two-way ANOVA, [K^+^]_o_ x IPI interaction, F_8,30_ = 7.66, *P* = 1.53 × 10^-5^, *n* = 3). In NAc, the effect of [K^+^]_o_ on PPR was less evident than in CPu, but showed an overall significance (Fig. 3D, Two-way ANOVA, main effect of [K^+^]_o,_ F_2,30_ = 15.15, *P* = 2.83 × 10^-5^, *n* = 3). Since we observed no effect of [K^+^]_o_ on 1p [DA]_o_, these effects on PPR were unrelated to initial P_r_ at either short (10 ms) or longer (100 ms) IPI (Fig. 3E, vertical slopes). These findings demonstrate directly that short-term plasticity of DA release can be dissociated from initial release, i.e. there is a release-insensitive short-term plasticity. Furthermore, they suggest that STD is governed by mechanisms governing axonal membrane polarisation/activation particularly in CPu.

**Figure 3.**
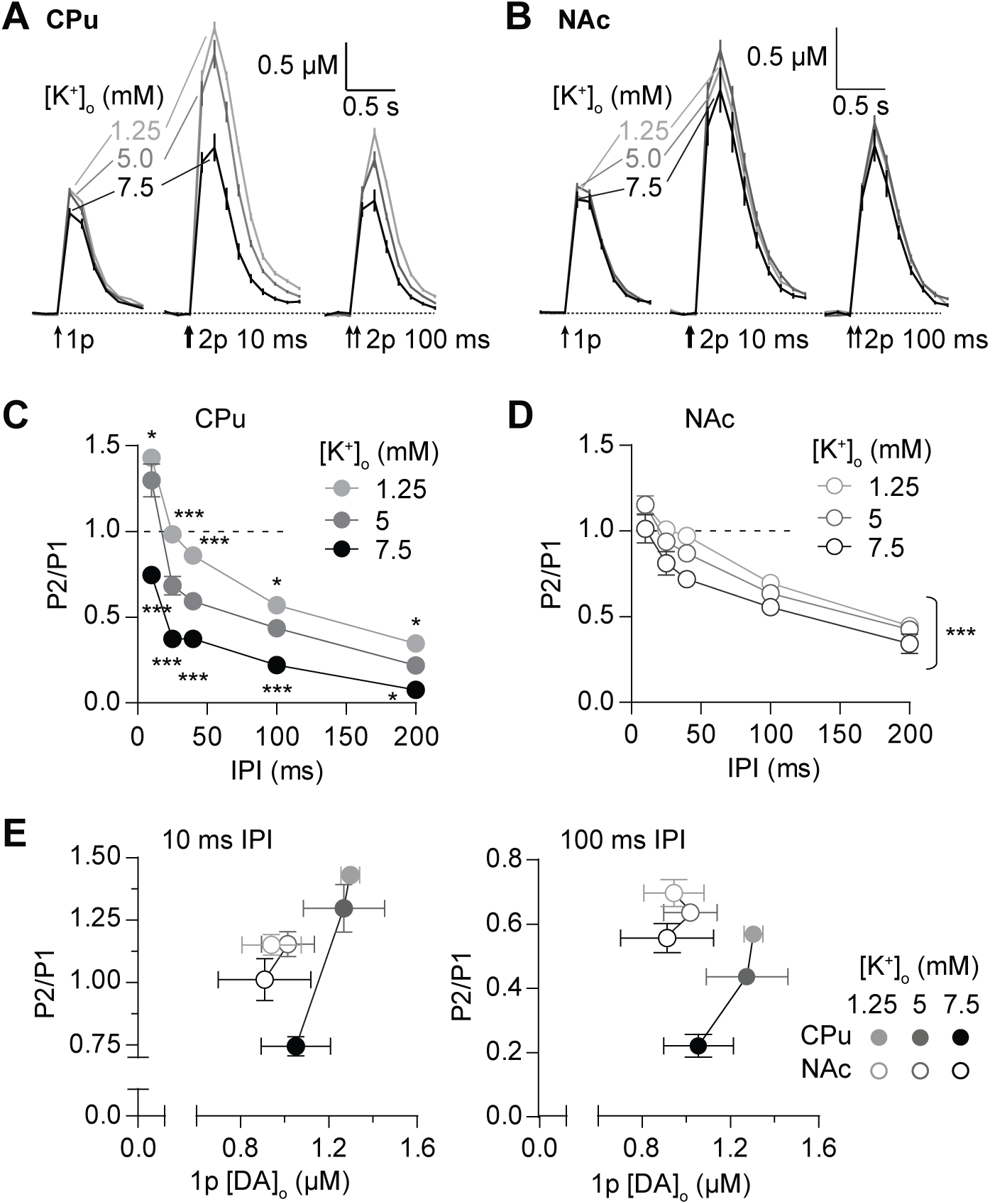
A release-insensitive mechanism of short-term depression is controlled by [K^+^]-dependent gating. **(A, B)** Mean profiles of [DA]_o_ (± SEM) elicited by single or paired electrical pulses in 1.25 mM (*light grey*), 5 mM (*dark grey*) and 7.5 mM [K^+^]_o_ (*black*) in CPu (A, *n* = 3) and NAc (B, *n* = 3). **(C, D)** Mean P2/P1 (± SEM) vs IPIs in 1.25 mM (*light grey*), 5 mM (*dark grey*) and 7.5 mM [K^+^]_o_ (*black*) in CPu (C) and NAc (D). **(E)** Mean P2/P1 (± SEM) vs. mean peak 1p [DA]_o_ (± SEM) in 1.25 mM (*light grey*), 5 mM (*dark grey*) and 7.5 mM (*black*) [K^+^]_o_ in CPu (*filled*) and NAc (*open circles*) at 10 ms IPI (*left*) and 100 ms IPI (*right*). Two-way ANOVA with Bonferroni’s test for *post-hoc* comparisons; \**P* < 0.05, \*\*\**P* < 0.001. Source data are provided as a Source Data file.

To identify whether the control of short-term plasticity by [K^+^]_o_ is accounted for by voltage-dependent effects, we tested whether the effects of [K^+^]_o_ on PPR could be substituted for, and prevented by broadly blocking K_v_ channels. We hypothesised that a K_v_ blocker (4-aminopyridine, 4-AP) should promote STD and prevent effects of changes in [K^+^]_o_. In CPu, broad inhibition of K_v_ channels by 4-AP (100 µM), unlike changes in [K^+^]_o_ alone, profoundly increased 1p [DA]_o_ (Fig. 4A, B, Two-way ANOVA, main effect of 4-AP: F_1,3_ = 33.63, *P* = 0.010, *n* = 4). This effect is consistent with the expected widening of the action potential by 4-AP [44] which will increase Ca^2+^ entry and neurotransmitter release for a single stimulus. In addition, 4-AP, similar to the effects of high [K^+^]_o_ (see Fig. 3), but unlike high [Ca^2+^]_o_ (see Fig. 2), reduced PPR at all IPIs, and also prevented the effects of changing [K^+^]_o_ on PPR (Fig. 4C, 4D, Two-way ANOVA, 4-AP x [K^+^]_o_ interaction, F_1, 12_ = 6.003, *P* = 0.031, *n* = 4). These findings indicate that inhibition of K_v_ channels promotes STD, and suggest that enhanced K_v_ currents relieve STD, and that hyperpolarizing/repolarizing conditions limit STD and promote subsequent release.

**Figure 4.**
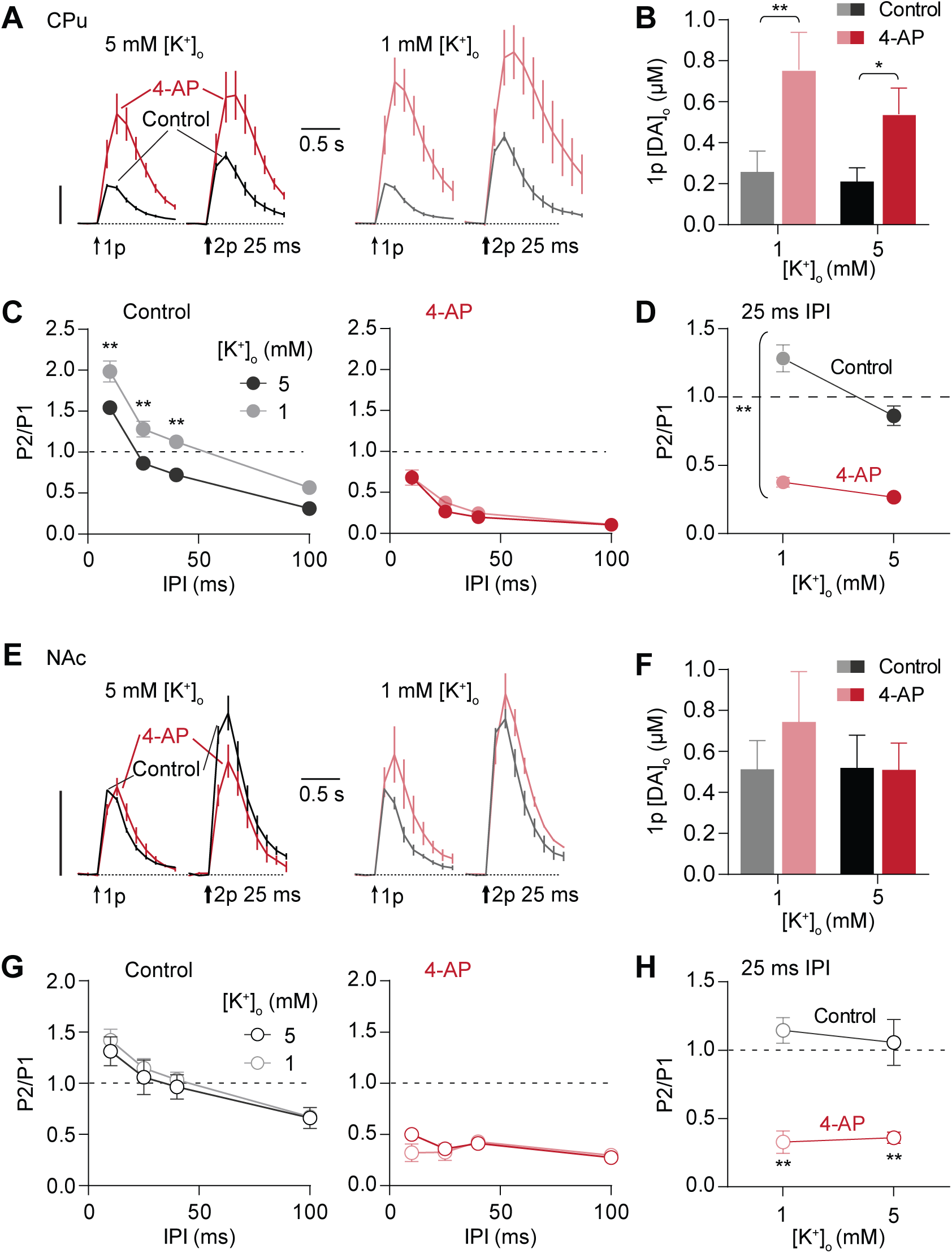
K^+^-dependent gating of short-term depression requires axonal Kv channels. **(A, E)** Mean profiles of [DA]_o_ transients (± SEM) in CPu elicited by single or paired electrical pulses in control conditions (*black*) or with 100 µM 4-AP (*grey*) in CPu (A, *n* = 4) and NAc (E, *n* = 3). Data are normalised to peak 1p-evoked [DA]_o_ (P1) in control conditions (*vertical scale bars*). **(B, F)** Mean 1p [DA]_o_ (± SEM) in CPu in 1 mM and 5 mM [K^+^]_o_ in control conditions (*black/grey*) or 4-AP (*reds*) in CPu (B) or in NAc (F). **(C, G)** Mean P2/P1 (± SEM) in CPu vs IPI in 5 mM (*black*) and 1 mM (*grey*) [K^+^]_o_ in control conditions (*left*) and 4-AP (*right*) in CPu (C) and in NAc (G). **(D, H)** Mean P2/P1 (± SEM) in CPu at 25 ms IPI in 1 mM and 5 mM [K^+^]_o_ in control conditions or 4-AP in CPu (D) and in NAc (H). (D), asterisks indicate interaction between [K^+^]_o_ and 4-AP. Two-way ANOVA with Bonferroni’s test for *post-hoc* comparisons; \**P* < 0.05, \*\**P* < 0.01. Source data are provided as a Source Data file.

In NAc, we observed no significant effect of 4-AP on 1p [DA]_o_ (Fig. 4E,F, Two-way ANOVA, *P* = 0.266, n = 3). However, 4-AP did decrease PPR at all IPIs, and there was no effect of varying [K^+^]_o_ in control conditions or in the presence of 4-AP (Fig. 4G, H). These findings verify that STD can be distinct from the effects on initial P_r_ and identify that STD is underpinned by the gating of axonal excitability.

### K^+^-dependent gating does not alter release-dependence of short-term plasticity

We investigated whether [K^+^]_o_-dependent STD could be a limiting factor that prevents short-term plasticity of DA release from responding to changes in P_r_ or whether yet further regulatory mechanisms might be indicated. We relieved STD (using low [K^+^]_o_) to test whether we could unmask a sensitivity of short-term plasticity to changes in initial release (using variation in [Ca^2+^]_o_). In CPu, 1p [DA]_o_ varied with [Ca^2+^]_o_ in a similar manner in 1 mM [K^+^]_o_ versus 5 mM [K^+^]_o_ (Fig. 5A-C) and although PPRs were elevated when [K^+^]_o_ was low (Fig. 5D) (as seen in Figs. 3C, 4C), there was no significant interaction between [K^+^]_o_ and [Ca^2+^]_o_ at IPIs of either 10 ms or 100 ms (Fig. 5D, Two-way ANOVA, 10 ms IPI: *P* = 0.086, 100 ms IPI: *P* = 0.654, n = 10). In NAc also, 1p [DA]_o_ varied with [Ca^2+^]_o_ in a manner that did not depend on [K^+^]_o_ (Fig. 5E-G) and there was no significant interaction between the effects of [K^+^]_o_ and [Ca^2+^]_o_ on PPR at either 10 ms or 100 ms IPI (Fig. 5H, Two-way ANOVA, 10 ms IPI: *P* = 0.963, 100 ms IPI: *P* = 0.883, n = 7). By limiting STD, we did not enhance Ca^2+^-dependent modulation of STF or STD in either CPu or NAc. Therefore, the mechanism through which [K^+^]_o_ modulates short-term plasticity is not masking or preventing short-term plasticity from being Ca^2+^-limited, in either CPu or NAc. This observation suggests that an additional mechanism may operate to limit the Ca^2+^- and release-dependence of short-term plasticity besides K_v_-regulation of axonal excitability and polarization.

**Figure 5.**
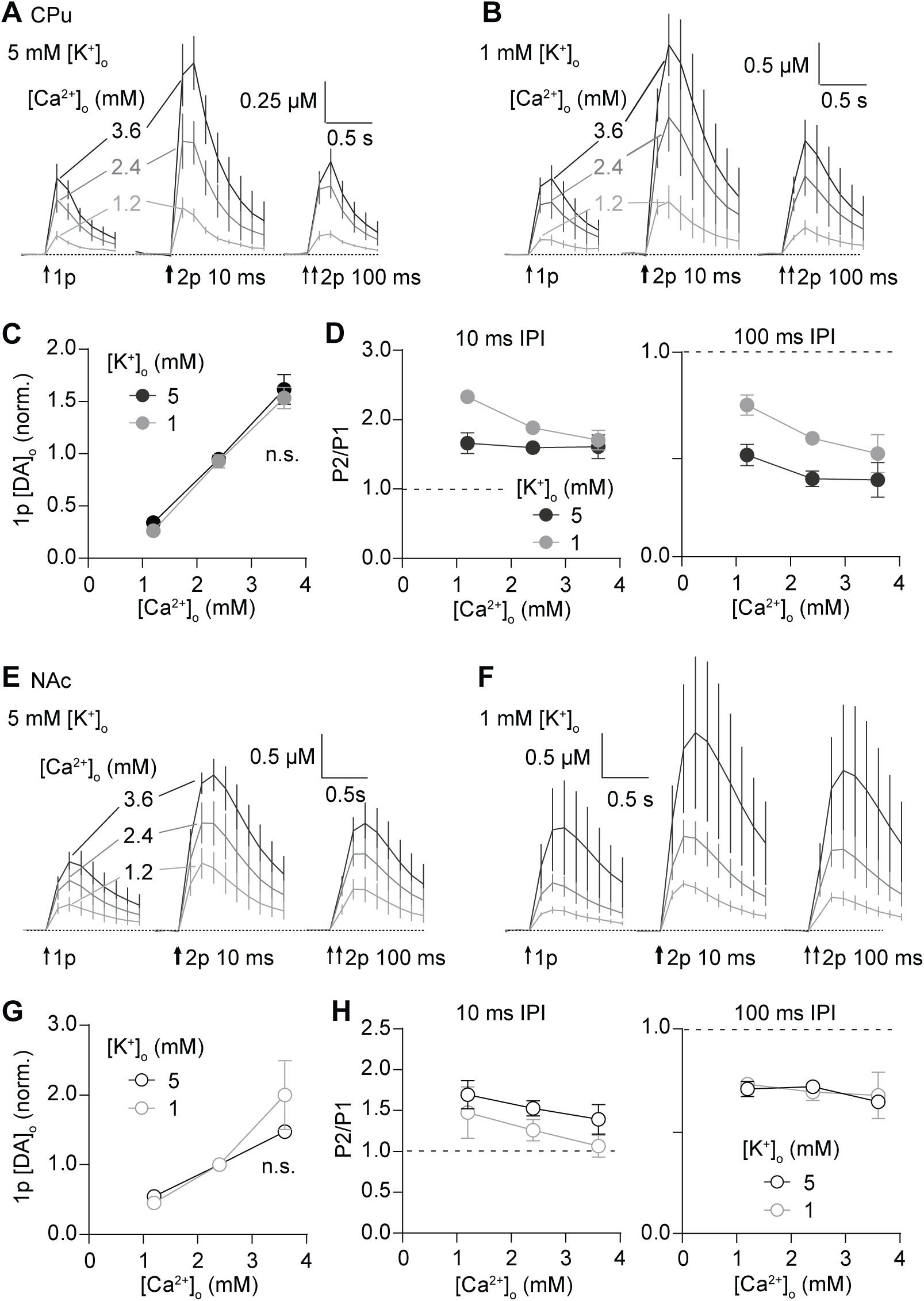
K^+^-dependent gating does not alter release-dependence of short-term plasticity. **(A,B,E,F)** Mean profiles of [DA]_o_ transients (± SEM) elicited by single or paired pulses in 1.2 mM [Ca^2+^] (*light grey*), 2.4 mM [Ca^2+^]_o_ (*dark grey*) and 3.6 mM [Ca^2+^]_o_ (*black*) in either 5 mM [K^+^]_o_ (*A,E*) or 1 mM [K^+^]_o_ (*B,F*) in CPu (A,B, *n* = 11) and in NAc (E,F, *n* = 7). **(C, G)** Mean peak 1p [DA]_o_ (± SEM, normalised to 2.4 mM [Ca^2+^]_o_) in 5 mM [K^+^]_o_ (*black*) and 1 mM [K^+^]_o_ (*grey*) in CPu (C) and in NAc (G). **(D, H)** Mean P2/P1 (± SEM) at 10 ms IPI (*left*) and 100 ms IPI (*right*) in 5 mM [K^+^]_o_ (*black*) and 1 mM [K^+^]_o_ (*grey*) in CPu (D) and in NAc (H). Two-way ANOVA with Bonferroni’s test for *post-hoc* comparisons. Source data are provided as a Source Data file.

### The dopamine transporter regulates short-term plasticity of dopamine release

Alongside its control of DA uptake, the dopamine transporter (DAT) has previously been shown to govern underlying DA release processes [29-32]. The DAT is also known to mediate electrogenic currents during DA binding and transport [27, 45] that have been shown to modulate the membrane potential of DA neurons *in vitro* [28]. We therefore investigated the hypothesis that DAT in striatum could consequently contribute to short-term plasticity.

To avoid adaptations to release seen after DAT knockout, we used uptake inhibitors to inhibit DAT function. We found that the monoamine uptake inhibitors cocaine and methylphenidate (MPH) and DAT inhibitor nomifensine significantly altered the pattern of short-term plasticity in a common manner. Cocaine (5 µM), MPH (5 µM) or nomifensine (10 µM) increased the peak amplitude and decreased the rate of decay of [DA]_o_ transients in CPu and NAc in keeping with DA uptake inhibition (Fig. 6A,C,G,I) and also prevented STF and significantly relieved STD (Cocaine: Fig. 6A,B, Two-way ANOVA, cocaine x IPI interaction, F_4, 30_ = 11.97, *P* = 6.24 × 10^-6^, *n* = 4, MPH: Fig. 6C,D, Two-way ANOVA, MPH x IPI interaction, F_4, 40_ = 5.316, *P* = 0.002, *n* = 5; Nomifensine: Fig. 6E,F, Two-way ANOVA, nomifensine x IPI interaction, F_4, 80_ = 21.12, *P* = 6.5 × 10^-12^, *n* = 3). These findings indicate that the DAT is a key regulator of short-term plasticity of DA release. At shortest IPI, DAT promotes STF, whereas at longer IPI, DAT clamps release to promote STD in CPu. In NAc, cocaine and MPH (we did not test nomifensine) prevented STF at 10 ms IPI (Fig. 6G,H, Two-way ANOVA, cocaine x IPI interaction, F_4, 30_ = 7.884, *P* = 1.82 × 10^-4^, *n* = 4; MPH: Fig 6I,J, Two-way ANOVA, MPH x IPI interaction, F_4, 20_ = 3.986, *P* = 0.016, *n* = 3) but in contrast to CPu, STD at longer intervals was not relieved. In NAc, the DAT regulated short-term plasticity at only short interpulse intervals. Since it is at these intervals that short-term plasticity is related to P_r_ (see Fig 1), these data suggest that it is the effect of DAT inhibition on promoting subsequent DA release that underlies its effects on short-term plasticity at short intervals in NAc.

**Figure 6.**
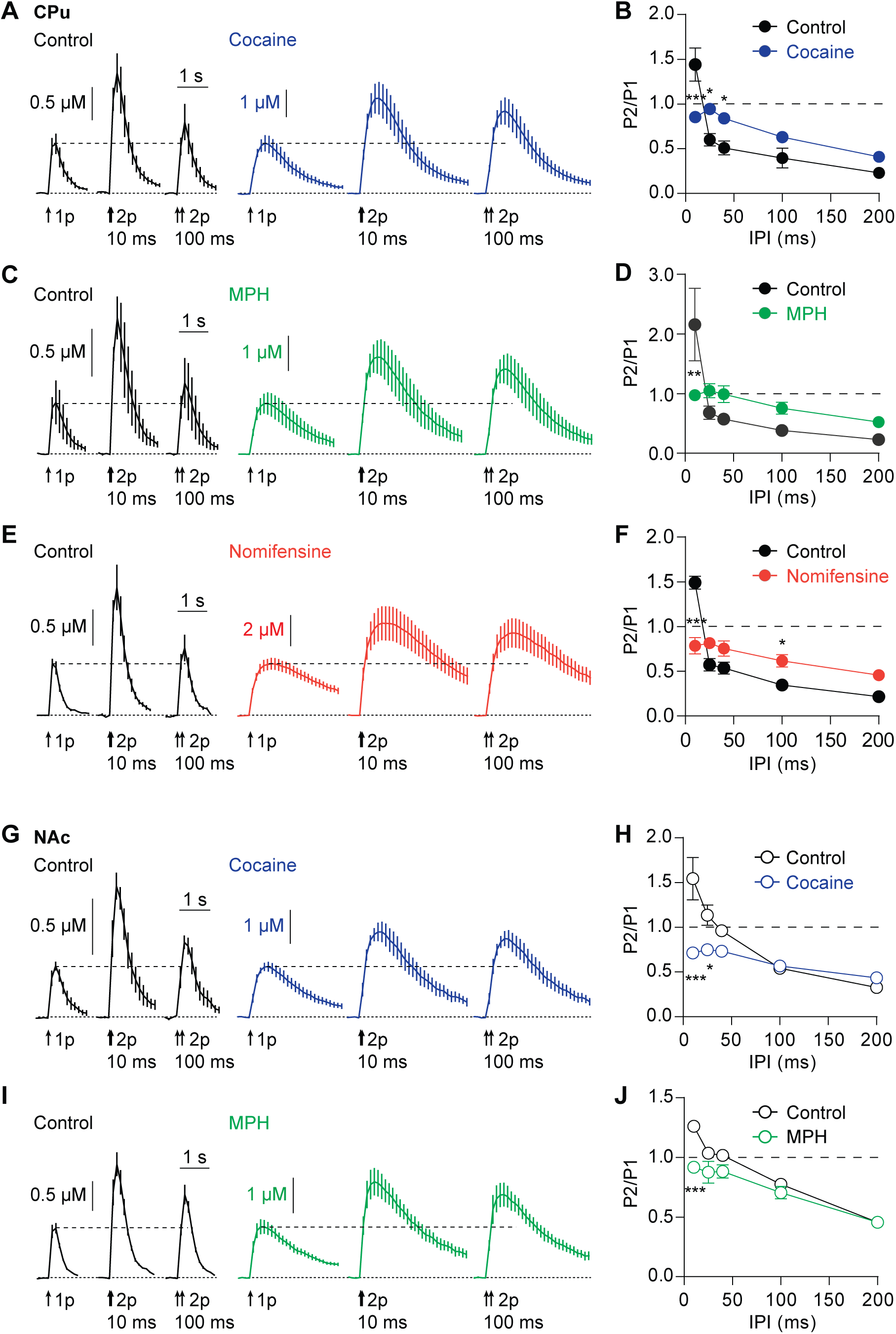
The dopamine transporter regulates short-term plasticity of dopamine release. **(A, G)** Mean profiles of [DA]_o_ transients (± SEM) elicited by single or paired electrical pulses in control conditions (*black*) or with 5 µM cocaine (*blue*) in CPu (A, *n* = 4) or NAc (G, *n* = 4). **(B, H)** Mean P2/P1 (± SEM) against IPI in control conditions (*black*) and with cocaine (*blue*) in CPu (B) or NAc (H). **(C, I)** Mean profiles of [DA]_o_ transients (± SEM) elicited by single or paired electrical pulses in control conditions (*black*) or with 5 µM methylphenidate (*MPH*, *grey*) in CPu (C, *n* = 5) or NAc (G, *n* = 3). **(D, J)** Mean P2/P1 (± SEM) vs IPI in control conditions (*black*) and with MPH (*green*) in CPu (D) or NAc (J). **(E)** Mean profiles of [DA]_o_ transients (± SEM) elicited by single or paired electrical pulses in control conditions (*black*) or with 10 µM nomifensine (*red*) in CPu (*n* = 3). **(F)** Mean P2/P1 (± SEM) vs IPI in control conditions (*black*) and with nomifensine (*red*) in CPu. Two-way ANOVA with Bonferroni’s test for *post-hoc* comparisons; \**P* < 0.05, \*\**P* < 0.01, \*\*\**P* < 0.001. Source data are provided as a Source Data file.

We ruled out activation of D_2_ receptors as contributing to DAT-mediated changes in short-term plasticity. At longer interpulse intervals of 2-3 seconds when sufficient time has elapsed for D2 receptors to have been activated, the D2 antagonist L-741626 in the presence of cocaine can enhance evoked [DA]_o_ at a second stimulus (**Suppl Fig. S3A,B**, Two-way ANOVA, main effect of L-741-626, F_1,64_ = 28.96, P < 0.0001, *n* = 4). However, the effect of cocaine on STF and STD at shorter intervals spanning 10-200 ms was not modified by the D2 antagonist (**Suppl. Fig S3C**, Two-way ANOVA, main effect of L-741,626, F_1,20_ = 2.76, P > 0.05, n = 4). We also ruled out potential local anaesthetic actions of cocaine on voltage-gated Na^+^ channels (VGSCs), since lidocaine (5 µM), an inhibitor of VGSCs, did not significantly alter short-term plasticity (**Supp Fig. S4**), consistent with reports that cocaine does not inhibit VGSCs at the concentration used in this study [46].

We tested whether the effects of DAT inhibition on short-term plasticity were dependent on synapsin III which has been suggested to mediate the role of the DAT in vesicle segregation and in limiting release [29]. However, the effects of cocaine on short-term plasticity persisted in mice lacking synapsin III (**Suppl. Fig. S5**), indicating that the DAT regulates short-term plasticity via a synapsin III-independent or other redundant mechanism.

### The DAT limits release-dependence of short-term plasticity in CPu

Since the DAT operates a clamp on DA release in CPu that drives STD, we tested whether the DAT might prevent short-term plasticity from being Ca^2+^- and release-dependent. We tested whether there was a stronger relationship between [Ca^2+^]_o_ and PPR in CPu, in the presence of cocaine. There was no effect on PPR of changing [Ca^2+^]_o_ in the absence of cocaine (Fig. 7A,B, Two-way ANOVA, *P* = 0.209, n = 4), but by contrast in the presence of cocaine there was a significant effect of [Ca^2+^]_o:_ PPR at IPIs of 10-40 ms were significantly elevated in 1.2 versus 2.4 mM [Ca^2+^]_o_ (Fig. 7A,C-E, Two-way ANOVA, [Ca^2+^]_o_ x IPI interaction, F_3, 24_ = 6.137, *P* = 0.003, *n* = 4). There was a significantly stronger effect of [Ca^2+^]_o_ on PPR in the presence of cocaine than in control conditions at 25 ms (Fig. 7D; Two-way ANOVA, cocaine x [Ca^2+^]_o_ interaction; F_1, 12_ = 7.86; *P* = 0.016; *n* = 4) and 40 ms IPI (Fig. 7D; Two-way ANOVA, cocaine x [Ca^2+^]_o_ interaction; F_1, 12_ = 4.72, *P* = 0.050, *n* = 4), but not at 10 ms or 100 ms IPI. These effects were not due to a potential electrochemical change in dopamine adsorption/desorption kinetics at the electrode that might occur with a change in divalent cations, since these effects of cocaine prevailed when the reduction in [Ca^2+^]_o_ was compensated by substitution with Mg^2+^ (**Suppl. Fig. S6**). Inhibition of the DAT therefore relieves a limiting clamp on a relationship between PPR and initial release, suggesting that the DAT limits the release-dependence of short-term plasticity and drives STD.

**Figure 7.**
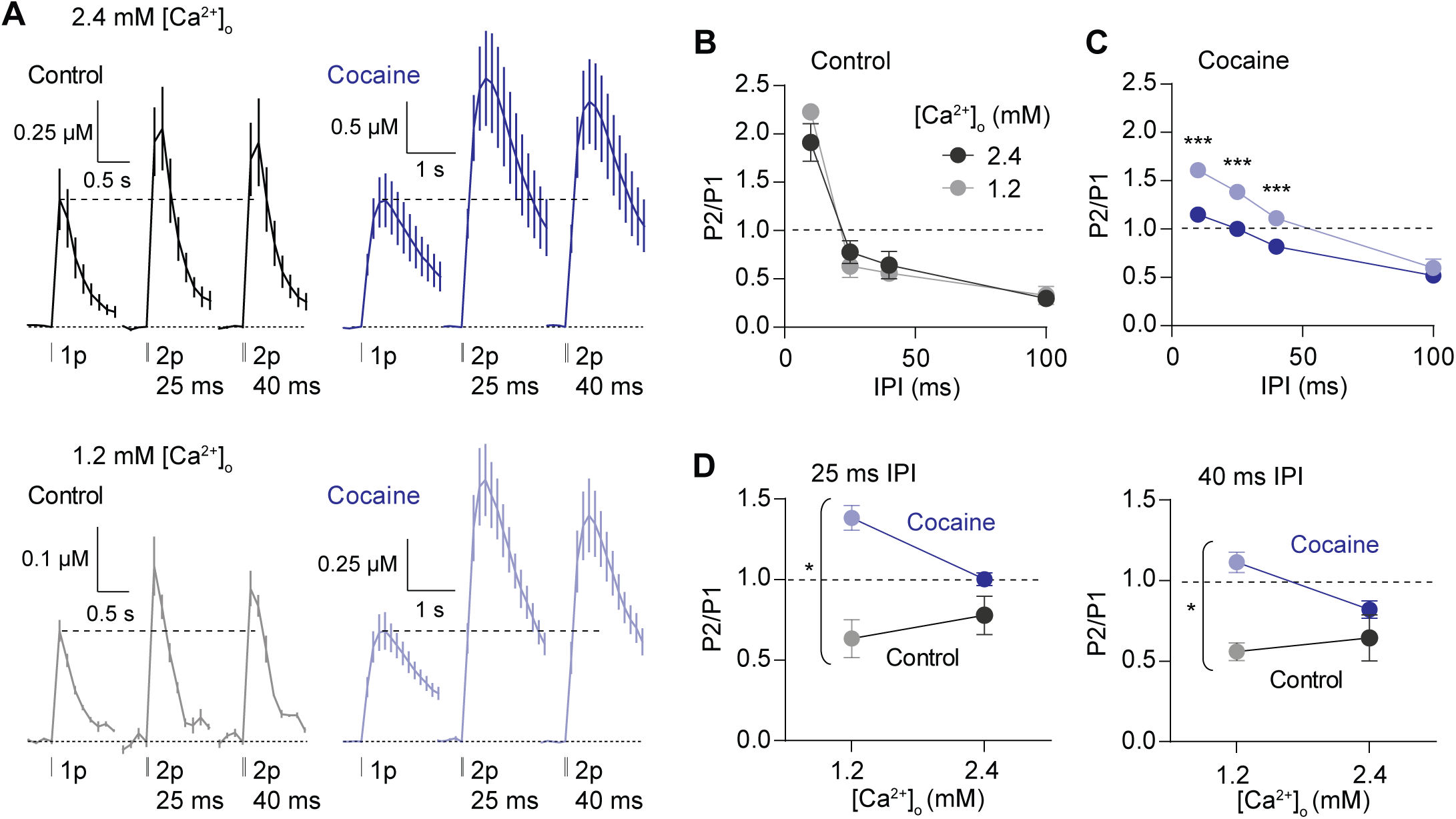
The DAT limits release-dependence of short-term plasticity in CPu. **(A)** Mean profiles of [DA]_o_ transients (± SEM, vertical scale normalised to 1p in 2.4 mM [Ca^2+^]_o_) (*n* = 4) elicited by single or paired electrical pulses in 2.4 mM (*upper*) and 1.2 mM [Ca^2+^]_o_ (*lower*) in control conditions (*black*) or 5 µM cocaine (*blue*). **(B, C)** Mean P2/P1 (± SEM) vs. IPI in 1.2 mM (*light*) and 2.4 mM [Ca^2+^]_o_ (*dark*) in control conditions (B) or cocaine (C). **(D)** Mean P2/P1 (± SEM) in 1.2 mM and 2.4 mM [Ca^2+^]_o_ in control conditions (*black-grey*) or cocaine (*blue*) at 25 ms IPI (*left*) or 40 ms IPI (*right*). Two-way ANOVA with Bonferroni’s test for *post-hoc* comparisons; \**P* < 0.05, \*\**P* < 0.01, \*\*\**P* < 0.001. Source data are provided as a Source Data file.

### The DAT maintains K^+^-dependent gating of short-term plasticity

We noted that the effect of DAT inhibitors on STD resembled the effect of low [K^+^]_o_. Since inhibition of the DAT might prevent DAT-mediated depolarising conductances and modify axonal hyperpolarisation/ repolarisation, we tested whether cocaine and low [K^+^]_o_ relieve STD via an overlapping mechanism. We identified whether cocaine precluded the effect of varying [K^+^]_o_ (1 vs 5 mM) on short-term plasticity, in CPu and NAc.

In CPu, in the absence of cocaine, PPR were elevated in lower [K^+^]_o_ (Fig. 8A,B, Two-way ANOVA, main effect [K^+^]_o_, F_1, 18_ = 20.47, *P* = 2.62 × 10^-4^, *n* = 4) as seen in Fig. 3. Cocaine increased 1p-evoked peak [DA]_o_ to a similar extent in 5 mM versus 1 mM [K^+^]_o_ (Fig. 8C) but prevented the effect on PPR of varying [K^+^]_o_ (Fig. 8D, Two-way ANOVA, [K^+^]_o_ x cocaine interaction, F_1, 12_ = 7.686, *P* = 0.017, *n* = 8). The limited ability of low [K^+^]_o_ to increase PPR was not due to a ceiling effect (such as DA depletion due to increased 1p release); cocaine also prevented the opposite effects of a rise in [K^+^]_o_ (to 7.5 mM) from decreasing PPR (**Suppl. Fig. S7**). In NAc, PPR were slightly elevated in lower [K^+^]_o_ (Fig. 8E,F, Two-way ANOVA, main effect [K^+^]_o_, F_1,12_ = 6.840, *P* = 0.023, *n* = 3). Cocaine increased 1p-evoked peak [DA]_o_ to a similar extent in 5 mM versus 1 mM [K^+^]_o_ (Fig. 8G), and, as in CPu, prevented the effect of varying [K^+^]_o_ on PPR (Fig. 8F,H, Two-way ANOVA, [K^+^]_o_ x cocaine interaction, F_1,8_ = 6.756, *P* = 0.032, *n* = 3). The effects of [K^+^]_o_ on short-term plasticity are therefore abolished by cocaine, suggesting that short-term plasticity is regulated by a pathway common to [K^+^]_o_ and DAT function. DAT function appears to regulate short-term plasticity by limiting Ca^2+^-dependent gating whilst supporting [K^+^]_o_-dependent modulation.

**Figure 8.**
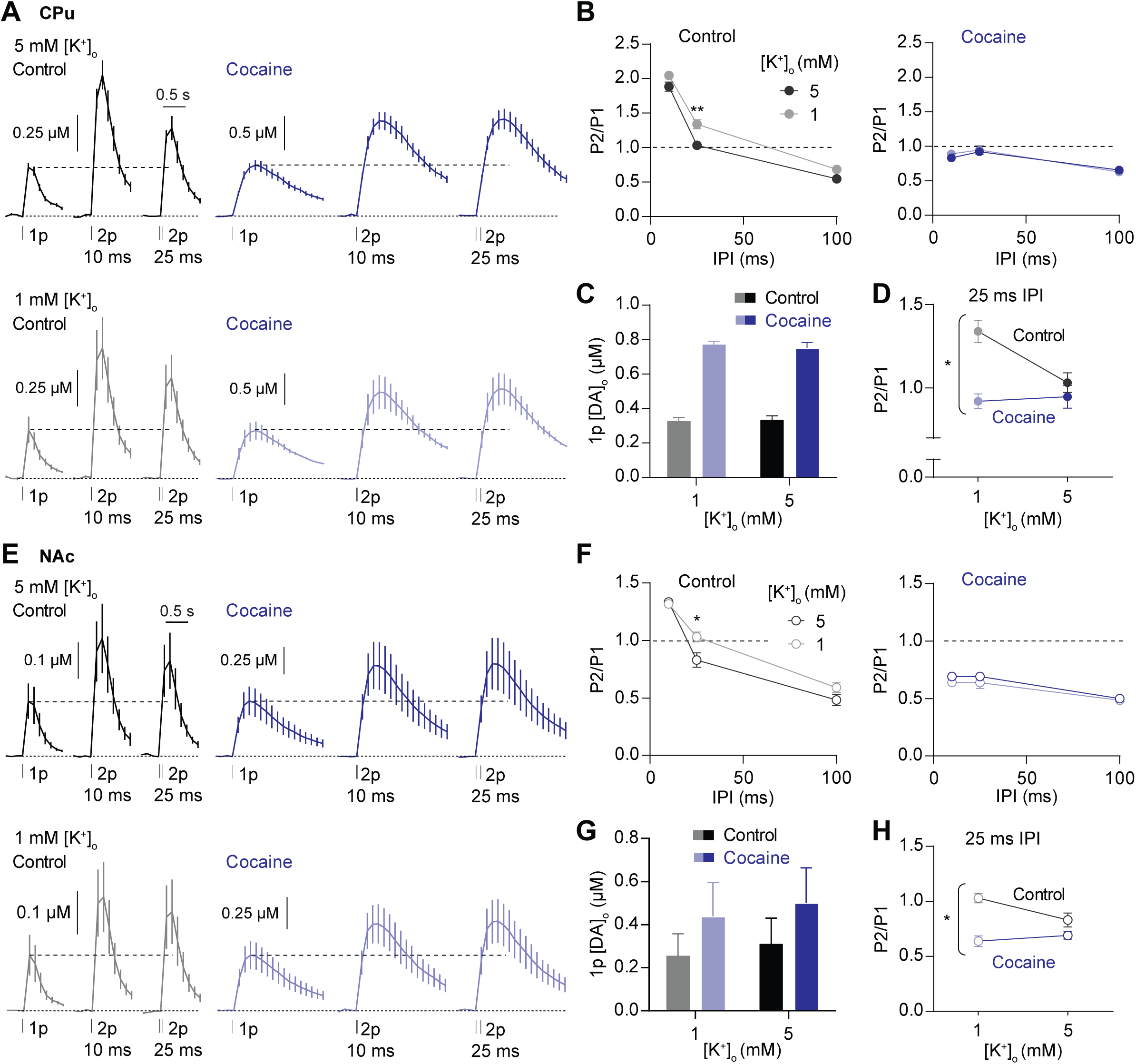
The DAT and [K^+^]_o_ gate short-term plasticity through overlapping mechanisms. **(A,E)** Mean profiles of [DA]_o_ transients (± SEM) elicited by single or paired electrical pulses in control conditions or cocaine in 5 mM [K^+^]_o_ (*upper*) and 1 mM [K^+^]_o_ (*lower*) in CPu (A, *n* = 4) and in NAc (E, *n* = 3). **(B, F)** Mean P2/P1 (± SEM) vs. IPI in 5 mM [K^+^]_o_ (*black*) and 1 mM [K^+^]_o_ (*grey*) in control conditions (*left*) or cocaine (*right*) in CPu (B) and in NAc (F). **(C, G)** Mean 1p [DA]_o_ (± SEM) in control conditions (*black-grey*) or cocaine (*blue*) in CPu (C) and in NAc (G). **(D, H)** Mean P2/P1 (± SEM) in CPu at 25 ms IPI in 1 mM and 5 mM [K^+^]_o_ incontrol conditions or with cocaine in CPu (D) and in NAc (H). Two-way ANOVA with Bonferroni’s test for *post-hoc* comparisons; \**P* < 0.05, \*\**P* < 0.01. Source data are provided as a Source Data file.

### The DAT and K^+^ regulate axonal activation

Finally, to validate that [K^+^]_o_ and the DAT are able to directly modulate the activation of DA axons, through mechanisms upstream of vesicular P_r_, we imaged Ca^2+^ dynamics in a population of DA axons in CPu using genetically encoded calcium indicator GCaMP6f expressed in DAT-Cre:Ai95D mice (Fig. 9A). We activated DA axons using the same single and paired stimulus pulses and conditions used to drive DA release, for two representative IPIs of 10 and 40 ms. We noted firstly, in control conditions for each experiment, that axonal Ca^2+^ levels evoked by 2p were significantly greater than those evoked by 1p, and were greater for IPIs of 10 ms than 40 ms (*P*<0.001, two-tailed paired t-test, n = 10) (Fig. 9B-D), which paralleled our observations of evoked [DA]_o_ throughout this study (e.g. see Fig. 1A,C). When we increased [K^+^]_o_ from 5 mM to 7.5 mM, Ca^2+^ levels in DA axons evoked by paired pulses (normalised to 1p) were slightly but significantly decreased for both IPI (Fig. 9B,C, Two-way ANOVA, main effect of [K^+^]_o_, F_1, 8_ = 25.95, *P* =0.0009, N = 5 animals), consistent with reduced re-activation of DA axons by a second pulse and the enhanced STD of DA release (see Fig. 3C). Furthermore, when we inhibited DATs with cocaine (5 µM), we found that Ca^2+^ levels evoked in DA axons by paired pulses were significantly modified, through a significant interaction between cocaine and IPI, which significantly increased Ca^2+^ levels at IPI of 40 ms, to levels seen at 10 ms (Fig 9D,E, Two-way ANOVA, interaction, F_1, 8_ = 23.19, *P* = 0.0013, N = 5 animals). This interaction broadly paralleled the impact of DAT inhibition on DA release at these IPIs (compare with Fig. 6A,B), and was consistent with a reduced STD in DA release being due at least in part to elevated re-activation of DA axons in the absence of DAT action.

**Figure 9.**
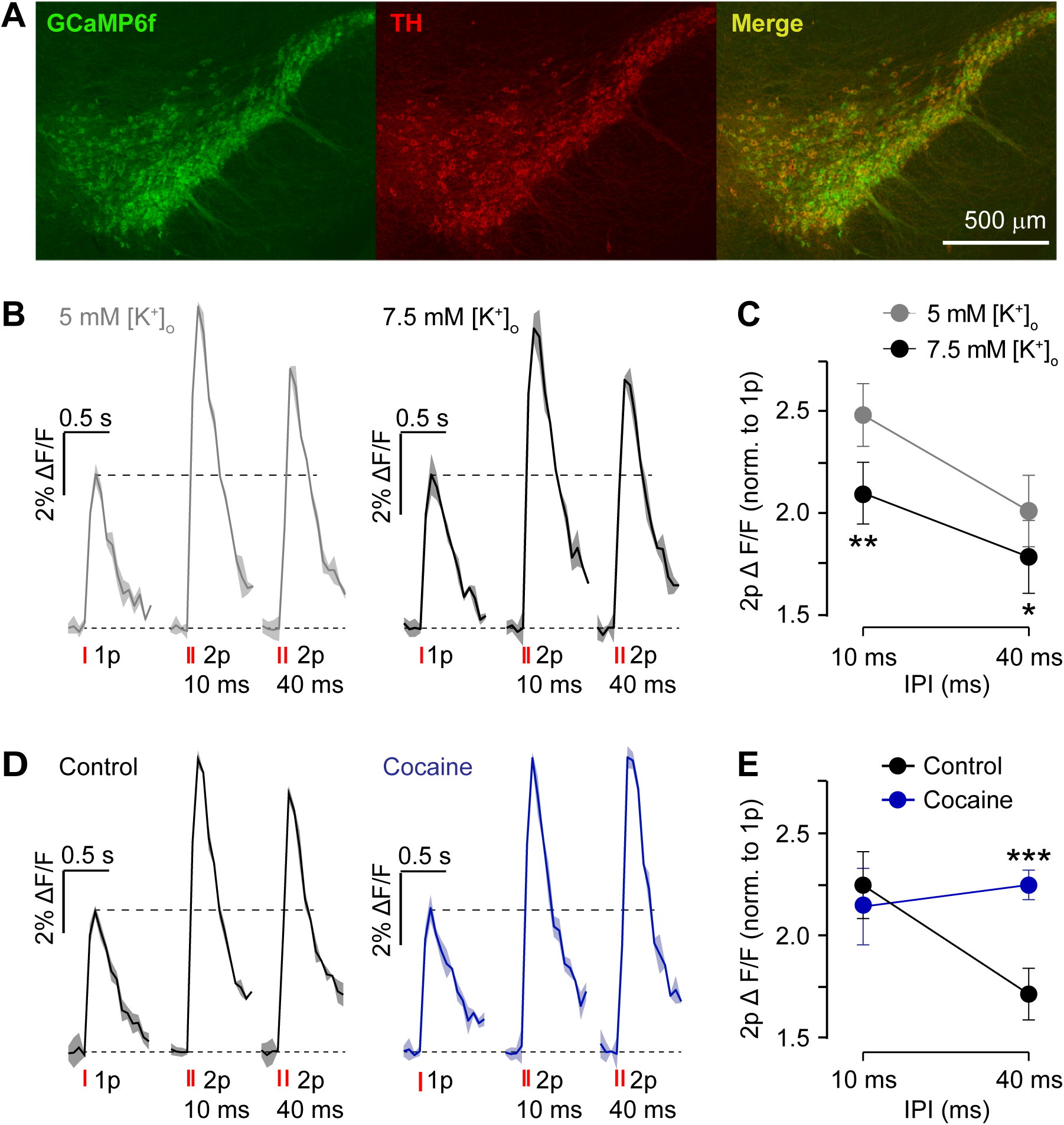
The DAT and [K^+^]_o_ gate intracellular Ca^2+^ dynamics during paired-pulse stimulations. **(A)** Images of VTA and SNc from DAT-Cre:Ai95D mice showing GCaMP6f eGFP expression (*green*) in TH-positive neurons (*red*). **(B,D)** Examples of Ca^2+^-imaging responses (changes to GCaMP6f fluorescence, ΔF/F) (mean ± SEM from duplicate) in DA axon population imaged in CPu in response to single or paired electrical pulses at IPI of 10 and 40 ms in (B) 5mM [K^+^]_o_ (*grey*) or 7.5 mM [K^+^]_o_ (*black*), or (D) in control conditions (*black*) or the presence of cocaine (5 µM) (*blue*). Data are rescaled to 1p in each condition to control for photobleaching. **(C,E)** Mean peak values (± SEM) for GCaMP6f ΔF/F evoked by 2 pulses vs. IPI. Data are normalised to value for 1 pulse (N = 5 animals). Two-way ANOVA with Fishers LSD test for *post-hoc* comparisons: \**P* < 0.05, \*\**P* < 0.01, \*\*\**P* < 0.001. DHβE (1 µM) is present throughout. Source data are provided as a Source Data file.

## Discussion

We addressed whether the mechanisms that control the short-term dynamic probability of DA release, in dorsal and ventral striatum, are governed by classic release-dependent or other release-independent mechanisms. We show that short-term plasticity is governed in only a limited manner by Ca^2+^-dependent regulation of release probability, which participates in determining STF but not STD, and to greater extent in ventral than dorsal striatum. We reveal that mechanisms insensitive to the initial level of release drive strong STD, which is therefore not limited by the vesicular pool. Rather, we find that axonal excitability and the DAT are major players in controlling STD, and they dominate over Ca^2+^-dependent gating. The DAT appears to be a master regulator that sets the dynamic level of DA release and its resulting STF, and clamps release leading to release-insensitive STD particularly in dorsal striatum. The differences seen between dorsal and ventral striatum could lead to divergent DA outputs in response to changes in action potential firing and to striatal modulatory inputs.

### Calcium-dependent facilitation

Studies of short-term plasticity at fast central synapses typically demonstrate an inverse relationship between PPR and initial P_r_; STF is observed when P_r_ is low, and STD when P_r_ is high [47]. Here, we show that, in NAc, an inverse relationship between STF and initial DA release could be observed for only short IPI corresponding to instantaneous frequencies seen during fast burst firing (≥25 Hz), but the STD seen at lower frequencies did not vary with initial release. By contrast, in CPu, there is limited relationship between Ca^2+^-limited initial release and the dynamic subsequent plasticity of DA release. Here, other release-independent mechanisms seem particularly influential. Given this overall apparent divergence from mechanisms typically operating at classic fast synapses, it is notable that some aspects of the molecular machinery supporting presynaptic active zones for DA release also diverge from that supporting fast synapses (e.g. RIM- and ELKS-dependence) [17].

STF at other synapses is often attributed to summation of residual Ca^2+^ [48, 49], or Ca^2+^-dependent ultrafast recruitment of vesicles [50]. Ca^2+^ dynamics can vary with local Ca^2+^ buffering mechanisms [51]. In NAc, we saw a more pronounced Ca^2+^-dependent STF than in CPu, despite a weaker relationship between [Ca^2+^]_o_ and initial DA release in NAc. These regional differences might correspond to differences in Ca^2+^ binding proteins. For example, the high-affinity fast Ca^2+^ buffer calbindin-D_28k_ is expressed at 2-3-fold greater levels in DA neurons of ventral tegmental area (VTA) which more strongly innervate NAc than neurons of substantia nigra pars compacta (SNc) which more strongly innervate CPu [52-54]. Calbindin might provide extra initial Ca^2+^ buffering capacity to limit initial release in NAc, but become saturated at subsequent stimuli, which in other neurons promotes STF [55, 56].

### Release-insensitive depression

At longer interpulse intervals (40-200 ms) corresponding to a range of physiological firing frequencies (5-25 Hz), there is strong STD of DA release. This is not due to D_2_ autoreceptor (D2R) activation at these intervals. STD of release of other transmitters can result from depletion of readily-releasable vesicles [57], but the STD in DA release does not seem to result from a limited availability of DA vesicles. Not only is a low fraction of presynaptic DA estimated to be released after stimulation [14,17,58,59], but moreover STD in the present study was not relieved by reducing initial release (by lowering [Ca^2+^]_o_). Ca^2+^-dependent inactivation of VGCCs by a Ca^2+^ sensor has also been proposed at some central synapses [60-62] but since low [Ca^2+^]_o_ did not relieve STD for DA release, this mechanism is unlikely to contribute here.

Since STD of DA release is not sensitive to the magnitude of initial release, it appears to be a release-insensitive depression. We found particularly in CPu, that STD varied with [K^+^]_o_, as seen at some other central synapses [63]. The variation in STD despite no change in initial release further demonstrate the uncoupling of STD from P_r_. The effects of [K^+^]_o_ suggested that factors underpinning the ability to re-activate DA axons, such as membrane polarity/excitability, are critical in determining STD for DA release. Using Ca^2+^ imaging, we validated that changes in [K^+^]_o_ modify the ability to activate DA axons at subsequent stimuli, consistent with axonal excitability being a critical factor in STD of DA release, upstream of local regulation of vesicular P_r_. A range of K_v_ channel classes can regulate axonal excitability and rates of repolarisation in CNS axons [64-67]. Varying [K^+^]_o_ would be expected to change the Nernstian driving force for active K^+^ currents [42], altering action potential waveform and repolarisation and therefore the degree of Na^+^ channel inactivation, or alternatively might alter the rate of K^+^ channel inactivation [43], leading to use-dependent changes in membrane potential which alter Na^+^ channel recruitment at a second pulse. We cannot distinguish here which of these opposing mechanisms dominates to govern short-term plasticity but in either scenario, Na^+^-channel recruitment would be altered.

The critical role for axonal excitability in DA STD is particularly pertinent given DA axon morphology. Midbrain DA neurons form small diameter, unmyelinated, extensively arborised axons each containing on the order of ∼10^4^ branch points [20, 68]. These morphological features will not readily favour reliable conduction of action potentials [69]. K^+^ or Na^+^ conductances are very influential in axonal conduction [63,70,71] and are likely to influence the fidelity or failure of action potential propagation through the DA axonal tree. Action potential properties and propagation failure could be key contributors to DA STD. As a consequence, the size of the DA axonal arbour invaded and the striatal field modulated by DA could be dynamically adapted by presynaptic activity and locally by neuromodulatory inputs. In this regard, it is noteworthy that nAChRs regulate DA release and can drive a strong STD that limits the frequency sensitivity in DA signals [11, 15]. ACh input to nAChRs might play a strategic role in shaping action potential propagation and fidelity throughout the axonal arbour.

Changes in propagation fidelity would be expected to have a greater impact on release where axonal arbours are more extensive. Notably, the axonal arbours formed by DA neurons projecting to CPu are more extensive than those projecting to NAc [19,20,72]. In parallel, short-term plasticity showed stronger dependence on K^+^-dependent gating in CPu than in NAc. This correlation suggests that action potentials in VTA DA neurons might more readily invade a fuller extent of their axonal tree than action potentials in SNc DA neurons, a speculation which could be tested in future studies with significant implications for signal processing by these neurons.

Together, these findings suggest that Ca^2+^-dependent mechanisms can modify amplitude of DA signals, but will not change the dynamic contrast in DA signals when DA neurons change their firing frequency, except at very highest frequencies in NAc. Conversely, mechanisms that modulate the driving forces on membrane potential and repolarisation, will particularly influence the dynamic contrast in DA signals.

### Role of the DAT in short-term plasticity

The DAT in striatum is known to regulate DA transmission through several means. DATs can curtail the extracellular summation and lifetime of [DA]_o_ through re-uptake [73], and can also limit the DA release process [29-32]. Correspondingly, we show firstly that DAT function promotes STF of DA release at short interpulse intervals (10 ms in CPu, and up to 25 ms in NAc) when short-term plasticity of DA release co-varies most strongly with initial release (see Fig. 1). But secondly, at longer intervals, corresponding to typical firing frequencies (5-25 Hz), when release is dominated by release-independent depression, we found that the DAT promotes STD, particularly in CPu. DAT function therefore can therefore promote both STF and STD. These effects are akin to changing both Ca^2+^-dependent P_r_ and K^+^-dependent excitability, but neither in isolation, and therefore indicate that the DAT is limiting initial DA release probability and also subsequent release through likely polarisation-dependent mechanisms.

The effects of DAT inhibition on STF at short IPIs seemed large given that, under control conditions, the relationship between initial release and STF was weak. This disparity is reconciled by our finding that DAT inhibition permitted Ca^2+^-dependent modulation of short-term plasticity, suggesting that the DAT is a critical player that controls and limits the relationship between Ca^2+^ and release probability. The mechanisms are not yet known. DATs have previously been suggested to inhibit Ca^2+^-dependent vesicle mobilisation via interactions with synapsins, with synapsin-3 indicated as a potential candidate [29, 30]. But our data with synapse-3 KOs do not readily support a synapsin3-dependent inhibition of vesicle recruitment by the DAT. The DAT is also an electrogenic transporter [74, 75] that mediates a depolarising current [27,28,45] and can interact directly with VGCCs [76] which might influence depolarization-dependent Ca^2+^ dynamics in DA axons. However, Ca^2+^ imaging in DA axons in CPu did not reveal an impact of DAT inhibition on Ca^2+^ levels for paired pulses at very short IPIs where STF occurs, suggesting that the impact of DATs on STF at very short intervals seems to be downstream of Ca^2+^ entry, e.g. in the mobilisation of vesicle pool or local regulation of vesicular P_r_. There might be redundancy within the synapsin family to continue to support a synapsin-dependent mechanism in the absence of synapsin3.

The DAT, like [K^+^]_o_, also acted to promote depression at longer, physiological IPIs (40 – 200 ms). The overlap between the effects on STD of the DAT, [K^+^]_o_ and K_v_ channel inhibitors suggested that the DAT acts through mechanisms that govern axonal activation. Using Ca^2+^ imaging we validated that DATs indeed limit the ability to activate DA axons at subsequent stimuli at these intervals. Thus, DATs might act to attenuate propagation of subsequent action potentials through the DA axon arbour. Since hyperpolarisation and low [K^+^]_o_ can promote action potential renewal and propagation in some axon types [77], the potential for the DAT, like high [K^+^]_o_, to depolarise membrane potential [27, 28] might contribute to poor axonal re-activation in DA axons. It is of note that DATs are widely distributed throughout the length of DA axons [78], and are found at locations thought to correspond to both release-active zones and inactive zones [17]. DAT function can also be modulated by DA D2 and D3-receptors [79, 80]. DATs could be ideally positioned to govern action potential propagation on axons and at branches, action potential waveform at release active zones (e.g. [66]) and/or the recruitment of vesicles or VGCCs at active zones, modulated by DA. The higher density of DA in dorsolateral striatum, and greater DAT transcript levels in SNc than VTA neurons [54], could lead to stronger limitations on action potential waveforms in CPu than NAc. Now defined, these hypotheses should be tested directly in future studies.

It is well known that DAT function can promote the frequency-dependence of DA release during ongoing activity [25,31,33,81,82]. These effects are consistent with the effects of uptake on extracellular summation between stimuli and also with the role of DAT in the dynamic probability of release we have identified here. The DAT might then determine both the fidelity and the spatial range of striatal DA transmission not only through its established roles in limiting DA diffusion, but also, by limiting the probability of DA release and activation of the axonal arbour.

In conclusion, we propose a hierarchy of intrinsic mechanisms that control short-term plasticity of DA release. Within this hierarchy, the DAT represents a ‘master regulator’, governing the balance between release-dependent and release-independent mechanisms that differently dominate in dorsal versus ventral striatum. In turn, DAT inhibitors such as cocaine will have profound effects on DA signalling through promoting P_r_, relieving STD, and altering the timecourse and spatial field of DA signals.

## Materials and Methods

### Animals

Experiments were carried out using adult male C57Bl6/J mice (Jackson Laboratories), heterozygous DAT^IRES^-^Cre^ mice, synapsin-III knockout (S3KO) mice, or DAT-Cre:Ai95D mice on a C57Bl6/J background. For experiments using optical stimulation of DA axons, male heterozygote DAT-internal ribosome entry site (IRES)-Cre mice were bred from homozygous DAT^IRES-Cre^ mice on a C57Bl6/J background (B6.SJL-Slc6^a3tm1.1(cre)Bkmn^/J, stock # 006660, Jackson Laboratories). S3KO mice were bred as described previously (Feng et al., 2002), and were kindly supplied by Professor HT Kao (Brown University). For Ca^2+^ imaging experiments, male heterozygous DAT-Cre:Ai95D mice (4 -8 weeks) were bred from homozygous DAT-Cre mice (B6.SJL-Slc6^a3tm1.1(cre)Bkmn^/J, JAX stock number 006660) crossed with homozygous Ai95D mice (B6;129S-Gt(ROSA)26Sor^tm95.1(CAG-GCaMP6f)Hze^/J, JAX stock number 028865). Animals were group-housed and maintained on a 12-hr light/dark cycle with *ad libitum* access to food and water. All procedures were performed in accordance with the Animals in Scientific Procedures Act 1986 (Amended 2012) with ethical approval from the University of Oxford, and under authority of a Project Licence granted by the UK Home Office.

### Surgery

Heterozygote DAT^IRES-Cre^ mice were injected intracerebrally with a Cre-inducible recombinant AAV serotype 5 vector containing an inverted gene for channelrhodopsin-2 fused in-frame with a gene encoding enhanced yellow fluorescent protein (pAAV-double floxed-hChR2(H134R)-EYFP-WPRE-pA), as we have used previously [11]. Mice were placed in a stereotaxic frame under isoflurane anaesthesia and a craniotomy was made above the injection site. Injections of 1 μL virus were given either unilaterally or bilaterally in either VTA (co-ordinates from Bregma in mm: AP -3.1, ML ±0.5, DV -4.4) or in the SNc (from Bregma in mm: AP -3.5, ML ±1.2, DV -4.0) using a 2.5 μL 33-gauge Hamilton syringe at 0.2 µL/min with a microinjector. The syringe was left in place for 10 minutes following each injection, then retracted slowly. Animals were maintained for at least 3 weeks following surgery to allow virus expression in striatum.

### Slice Preparation

Striatal slices were prepared as described previously [83, 84]. Mice were sacrificed by cervical dislocation and the brains removed and transferred to ice-cold HEPES-based buffer containing in mM: 120 NaCl, 20 NaHCO_3_, 6.7 HEPES acid, 5 KCl, 3.3 HEPES salt, 2 CaCl_2_, 2 MgSO_4_, 1.2 KH_2_PO_4_, 10 glucose, saturated with 95%O_2_/5%CO_2_. Acute 300 µm thick coronal striatal slices, containing both dorsal striatum (CPu) and nucleus accumbens core (NAc) were prepared in ice-cold HEPES-based buffer and cut using a vibratome (VT1000S or VT1200S; Leica). Slices were kept at room temperature in HEPES-based buffer for 1 hour before being transferred to the recording chamber and superfused at 1.8-2.0 ml/min in bicarbonate buffer-based artificial CSF (aCSF) containing in mM: 124.3 NaCl, 26 NaHCO_3_, 3.8 KCl, 2.4 CaCl_2_, 1.3 MgSO_4_, 1.2 KH_2_PO_4_, 10 glucose, saturated with 95% O_2_/5% CO_2_, at 31-33°C. Recording medium also contained dihydro-β-erythroidine (DHβE, 1 µM), an antagonist of β2 subunit-containing nAChRs. Slices were allowed to equilibrate for 30 minutes prior to recording.

### Voltammetry and Stimulation

Evoked extracellular DA concentration ([DA]_o_) was measured using FCV at carbon-fibre microelectrodes (fibre diameter 7-10 µm, tip length 50-100 µm) implanted to a constant depth of 100 µm. A triangular voltage waveform was scanned across the microelectrode (-700 to +1300 mV and back vs Ag/AgCl reference, scan rate 800 V/s) using a Millar voltammeter (Julian Millar, Barts and the London School of Medicine and Dentistry), with a sweep frequency of 8 Hz, as described previously [83]. This sampling rate is sufficient to capture the rising and falling phase of the DA transients; faster sampling rates do not change the data interpretations (not illustrated). Evoked currents were confirmed as DA by comparison of the voltammogram with that produced during calibration with applied DA in aCSF (oxidation peak +500-600 mV and reduction peak -200 mV). Currents at the oxidation peak potential were measured from the baseline of each voltammogram and plotted against time to provide profiles of [DA]_o_ versus time. Electrodes responded linearly to [DA]_o_ over the concentration range detected. Electrodes were calibrated after use in 2 µM DA in each experimental solution used, including all solutions where [Ca^2+^] was varied, since electrode sensitivity varies with divalent ion concentration [85]. Calibration solutions were made up immediately before use from stock solution of 2.5 mM DA in 0.1 M HClO_4_ stored at 4°C. Electrode sensitivities ranged from 5-25 nA/µM.

For experiments using electrical stimulation, DA release was evoked using a surface bipolar concentric Pt/Ir electrode (25 µm diameter, FHC) placed approximately 100 µm from the recording electrode. Stimulation pulses of 200 µs duration were applied at 0.6 mA unless described otherwise.

For optogenetic stimulation, DA release was evoked in striata from DAT^IRES-Cre^ mice conditionally expressing ChR2 in DA axons, using 2 ms full-field illumination from an LED system emitting light at 470 nm wavelength (OptoLED, Cairn Research) as described previously [11]. The LED system illuminated an area of 2.2 mm diameter. The current delivered by the LED power supply was set to produce a perimaximal light intensity (i.e. the minimum light intensity able to evoke maximum [DA]_o_ following a single 2 ms pulse). Since the perimaximal light intensity is dependent on the level of ChR2 expression, which varies between animals, the appropriate current was determined at the beginning of each experiment. Electrical or optical stimulations were delivered every 2.5 minutes, after which dopamine release was reproducible. Before acquisition of experimental data, peak evoked [DA]_o_ levels were allowed to reach this reproducible stable level.

All experiments were conducted in the presence of nAChR antagonist DHβE (1 µM) to prevent confounding effects of striatal ACh. Striatal nAChRs operate a profound control over striatal DA release: ACh released by striatal stimulation can directly drive axonal DA release [11] and limit dependence of DA release on presynaptic frequency [15, 83] which can mask other mechanisms operating presynaptically on DA axons. Inclusion of nAChR antagonists during local stimulation also makes evoked DA release in *ex vivo* slice sensitive to frequency of stimulation [15, 83] as seen *in vivo* after midbrain or medial forebrain bundle stimulation [81, 86].

### DA release study design and analyses

The term release probability P_r_ used for DA here is a composite measure of synaptic P_r_ (a function of vesicular P_r_ and the size of the pool of vesicles) and the number of release sites and fibres recruited by the stimulus. Short-term plasticity in DA release was explored by applying alternating single pulses (1p) or paired pulses (2p) with inter-pulse intervals (IPIs) of 10 ms – 200 ms in pseudorandom order and in triplicate at each recording site. IPIs of 40 to 200 ms fall within the range commonly observed during burst firing in DA neurons *in vivo* [3, 4]. IPIs of 10 to 25 ms have been observed during burst firing in rat DA neurons [4] but are particularly useful to interrogate short-term facilitation which occurs on this timescale [15, 25].

We calculated paired-pulse ratio (PPR) as a measure of short-term plasticity. We define PPR as the ratio P2/P1, where P1 is peak [DA]_o_ detected following 1p stimulus and P2 is the peak [DA]_o_ attributable to the second stimulation only. P2 was determined by subtracting the entire [DA]_o_ transient including decay phase after a single pulse from the summated paired-pulse response, and we therefore account for summation and decay resulting from uptake kinetics. Any enhanced spillover resulting from uptake inhibition will occur for single and paired pulses and should therefore be controlled for. We cannot control for variable fiber recruitment for different stimuli, but can control for variable modulation by other local inputs: PPR on the timecourses explored here is not differently modified at different intervals by D2 dopamine receptors (see Results), or by GABA receptors (Lopes and Cragg, unpublished observations). We use the term “release-dependent” plasticity to indicate a relationship between PPR and P1 [DA]_o_. “Release-insensitive” or “release-independent” refers to a PPR that varies independently of P1, or does not vary with P1.

Calcium concentration ([Ca^2+^]_o_) in aCSF was varied by varying [CaCl_2_]. In some experiments where stated, a reduction in CaCl_2_ was substituted with MgCl_2_. However, Ca^2+^/Mg^2+^ substitution was not routine because other effects of [Mg^2+^]_o_ can confound interpretation. Changes in [Mg^2+^] have been shown to alter Ca^2+^ currents through VGCCs [87, 88], alter G-protein-coupled receptor function, including dopamine D2 receptors [89], change the affinity of calcium-binding protein calbindin for Ca^2+^ [90], trigger complexing of Ca^2+^ by ATP [91], and alter the sensitivity of FCV electrodes to DA [85], and we find also that Ca^2+^/Mg^2+^ substitution reduces DA release to levels below those of reducing Ca^2+^ alone (**Supplementary Figure S6D**).

Where extracellular potassium concentration ([K^+^]_o_) was varied (1.0 - 7.5 mM), changes in osmolarity were corrected by varying [Na^+^]. The Na^+^- and K^+^-containing salts present in aCSF were varied as follows (in mM): In experiments with 1 mM [K^+^]_o_, salts were 127.1 NaCl, 1.0 KCl, 0 KH_2_PO_4_, 1.2 NaH_2_PO_4_; For 1.25 mM [K^+^]_o_ salts were 128.0 NaCl, 0.05 KCl, 1.2 KH_2_PO_4_; For 7.5 mM [K^+^]_o_, salts were 121.8 NaCl, 6.3 KCl and 1.2 KH_2_PO_4_. Recordings in control conditions were collected before and after each experimental manipulation at each recording site. Slices were equilibrated in each solution for at least 20 minutes before data were included for analysis. Experiments were carried out in either the dorsal half of the caudate-putamen (CPu) (dorsolateral and dorsomedial striatum) or nucleus accumbens core (NAc) (within 200 µm of the anterior commissure), one site per slice. Release is TTX-sensitive [11]. Manipulations of [Ca^2+^]_o_ or [K^+^]_o_ did not change DA decay rates.

Data were acquired and analysed using Axoscope 10.2, 10.5 (Molecular Devices) or Strathclyde Whole Cell Program (University of Strathclyde, Glasgow, UK) and locally-written scripts in Visual Basic for Applications (Microsoft). Data are expressed as mean ± standard error (SEM) and *n* = number of animals (the number of different biological replicates). Each experiment was performed at a single recording site in one brain slice. Within a given experiment at each single recording site, technical replicates were usually obtained in at least triplicate before averaging to obtain the value for that experiment at that recording site. Statistical comparisons were carried out using GraphPad Prism 6, 7 and 8 (GraphPad Software Inc.) using ANOVAs, post-hoc two-tailed t-tests with n = number of animals. Data were tested for normality before using parametric tests.

### Ca^2+^ imaging

An Olympus BX51Wl microscope equipped with a CAIRN Research OptoLED Lite system, Prime Scientific CMOS (sCMOS) Camera (Teledyne Photometrics), and a 40x/0.8 NA water-objective (Olympus) was used for wide-field fluorescence imaging of GCaMP6f in dopaminergic axons in dorsolateral CPu in *ex vivo* slices in response to single and paired electrical stimulus pulses. Images were acquired at 16.6 Hz frame rate every 2.5 minutes using Micro-Manager 1.4, with stimulation and recording synchronised using custom-written procedures in Igor Pro 6 (WaveMetrics) and an ITC-18 A/D board (Instrutech). Image files were analysed with Matlab R2017b and Fiji 1.5. We extracted fluorescence intensity from the region of interest (ROI) and from an equal background area where there was no GCaMP6f expression (on the stimulating electrode). After background subtraction, the Ca^2+^ transients were bleach-corrected by fitting an exponential curve function through both the baseline (2 s prior to stimulation) and the last 1 s in a 7.2 s recording window. Data are expressed as ΔF/F where F is the fitted curve. The order of single and paired stimulation pulses was alternated and equally distributed, and data were collected in duplicate before and after a change in extracellular experimental condition. Data are expressed as mean ± standard error of the mean (SEM) for each stimulus condition. N value is the number of animals. Statistical analyses used GraphPad Prism 7.03, for two-way ANOVA with post-hoc t-tests. All experiments were conducted in the presence of DHβE (1 µM). GCaMP6f responses were quantified as the mean ΔF/F value.

### Immunocytochemistry

We verified the specificity of GCaMP6f expression to dopaminergic structures by comparing direct eGFP fluorescence to immunoreactivity to tyrosine hydroxylase (TH-ir). Acute slices of midbrain and striatum were fixed overnight at 4 °C in 4% paraformaldehyde dissolved in PBS, then stored in PBS. After resectioning to 40 µm, free-floating sections were washed in PBS 5 × 5 min and incubated in 0.5% Triton X-100, 10% normal goat serum and 10% fetal bovine serum for 30 min. Slices were subsequently incubated overnight with 1:2000 primary (rabbit anti-TH; Sigma) antibody dissolved in PBS containing 0.5% Triton X-100, 1% normal goat serum and 1% fetal bovine serum. Sections were then washed with PBS 5 × 5 min and incubated for 2 hr at room temperature with 1:1000 secondary (DyLight 594 goat anti-rabbit; Jackson) antibody dissolved in PBS containing 0.5% Triton X-100, 1% normal goat serum and 1% fetal bovine serum. Sections were washed with PBS and mounted on gelled slides with Vectashield mounting medium (Vector Labs) and imaged using a Zeiss LSM880 (confocal) running Zen black version 2.3, at 20x, N.A. 0.8. Maximum intensity projection from a z-stack of height 30 µm was captured individually and the stack of the pictures were compressed. TH (red) was captured at 638-759 nm with 633 nm excitation light. GCaMP (green) was excited with 488 nm and captured at 493-630 nm.

### Drugs

DHβE and lidocaine were obtained from Tocris Biosciences or Ascent Scientific. Cocaine hydrochloride and nomifensine maleate were obtained from Sigma. Methylphenidate (Ritalin™) was obtained from Novartis. All drugs were dissolved in either de-ionised water, aqueous acid (nomifensine maleate) or ethanol (lidocaine) to make stock aliquots at 1000-10,000x final concentration and stored at -20 °C prior to use. DA uptake inhibitors were used at standard concentrations that have detectable outcomes on uptake kinetics concentrations but are lower than those that lead to run-down of release and/or have non-selective effects as local anaesthetics or at nAChRs [46,92,93].

## Acknowledgements

Funding sources: MRC, BBSRC Interdisciplinary Doctoral Training Program, Clarendon Fund, Monument Trust Parkinson’s UK Discovery Award, and Christ Church Oxford.

## Author contributions

M.D.C. designed and performed experiments, analysed and interpreted the data, and co-wrote the manuscript. N.J.P. designed and performed experiments, and analysed and interpreted the data. B.M.R, S.V.M. and M.Y.T performed experiments and analysed data. K.R.B., S.T. and S.J.C. supervised the work. S.J.C. designed the experiments, interpreted the data, and co-wrote the manuscript.

## Data Availability

The authors declare that all data sets generated and analysed during this study are available within this paper and its supplementary files. The source data underlying all figures and supplementary figures are provided as a Source Data file.

