## Supplemental Figures S1-S7 for "Plasticity in striatal dopamine release is governed by release-independent depression and the dopamine transporter"

### Supporting Data

### Figures S1-S7 and Legends

Source data are provided separately as a Source Data file.

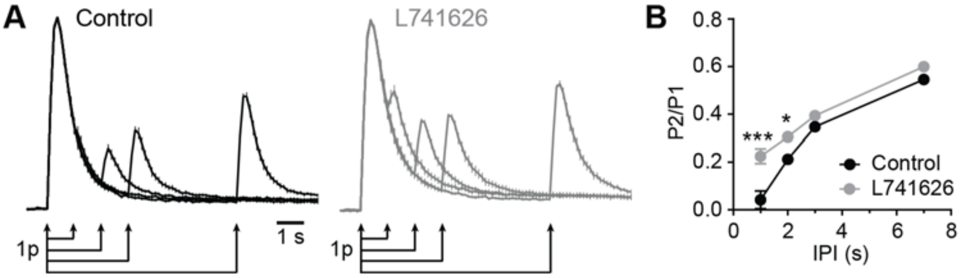


**Figure S1. D2 receptor antagonist L-741626 relieves STD at interpulse intervals of 1-2 seconds. (A)** Mean profiles of $[DA]_o$ transients (± SEM) evoked by pairs of single electrical pulses (*arrows*) in control conditions (*left*) and in the presence of $D_2$-receptor antagonist L741626 (1 µM, *right*) in CPu ($n$ = 3). Data are normalised to P1 peak $[DA]_o$ in each condition. **(B)** Mean P2/P1 ± SEM vs. IPI. At IPIs of 1-2 s, L741626 significantly increased P2/P1 (Two-Way ANOVA, effect of L741626, $F_{1,64}$= 36.27, $p$ < 0.0001, post-hoc Sidak's multiple comparison t-tests, ***$p$ < 0.001, *$p$ < 0.05). P2 values are obtained from the peak values of $[DA]_o$ seen at P2 after subtraction of $[DA]_o$ transients evoked by 1p from those evoked by 2p.

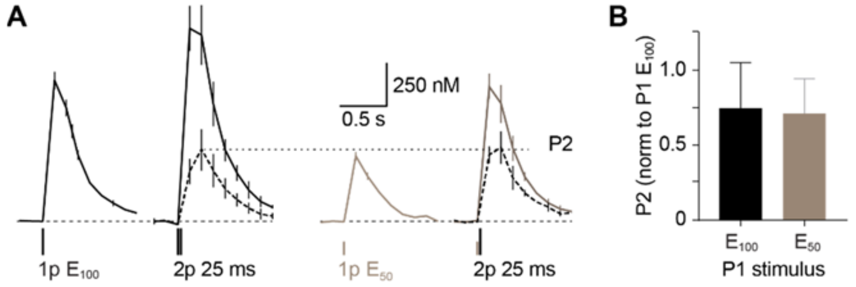


**Figure S2. Release at paired pulse is not determined by magnitude of initial evoked $[DA]_o$. (A)** Mean profiles of $[DA]_o$ transients (± SEM) evoked by 1 and 2 electrical pulses in CPu at an IPI of 25 ms after a first pulse at a maximal ($E_{100}$) or half maximal current ($E_{50}$). $[DA]_o$ evoked by the second stimulus (P2) ($E_{100}$ in both cases) evoked similar $[DA]_o$ after each initial stimulus. **(B)** Mean peak $[DA]_o$ (± SEM) evoked at P2 (normalised to P1 release with $E_{100}$) when did not vary when P1 was evoked by $E_{100}$ or $E_{50}$. Paired t-test: $p$ = 0.6638, N = 4.

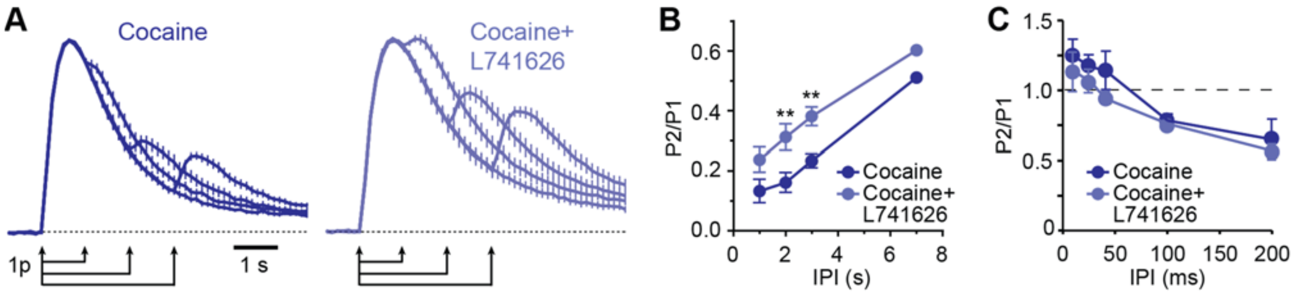


**Figure S3. D2 receptors do not modify short-term plasticity seen at short interpulse intervals (<200 ms) in CPu in the presence of cocaine. (A)** Mean profiles of $[DA]_o$ transients (± SEM) evoked by pairs of single electrical pulses (*arrows*) in the presence of cocaine (5 µM, *left*) and cocaine plus D2-receptor antagonist L741626 (1 µM, *right*) in CPu (*n* = 4). Data are normalised to 1p-evoked $[DA]_o$ (P1) in each condition. **(B)** Mean P2/P1 ± SEM vs. IPI in cocaine only (*dark blue*) or cocaine plus L741626 (*light blue*). At longer IPIs ≥ 1s, L741626 significantly increased P2/P1 in the presence of cocaine vs cocaine only (Two-Way ANOVA, effect of L741626, $F_{1,64}$= 28.96, $p < 0.0001$, post-hoc Sidak's multiple comparison t-tests, $^{**}p < 0.01$). P2 are the peak values of $[DA]_o$ seen after subtraction of $[DA]_o$ transients evoked by 1p from those evoked by 2p. **(C)** Mean P2/P1 ± SEM vs. IPI in cocaine only (*dark blue*) or cocaine plus L741626 (*light blue*) (*n* = 4). At short IPIs ≤ 200 ms, L741626 did not modify P2/P1 (Two-Way ANOVA, effect of L741626, $F_{1,20}$=2.76, $p > 0.05$). DHβE is present throughout.

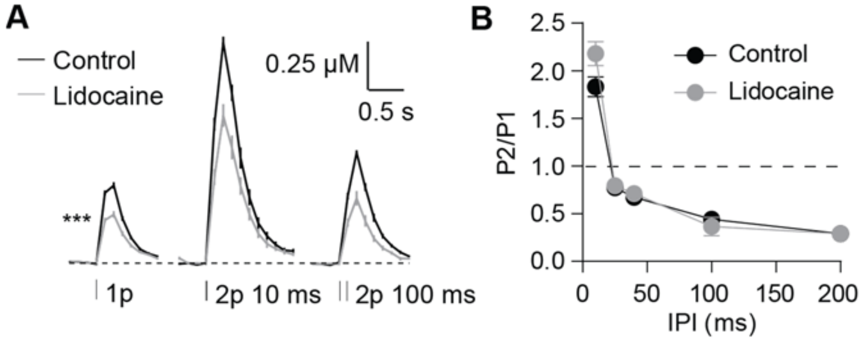


**Figure S4. Voltage-gated Na$^+$ channel inhibitor lidocaine does not replicate the effects of cocaine on short-term plasticity in CPu. (A)** Mean profiles of $[DA]_o$ transients (± SEM) evoked by electrical stimulation in control conditions (*black*) or in the presence of 5 µM lidocaine (*grey*) in CPu (*n* = 3). Lidocaine significantly decreased 1p $[DA]_o$ ($p < 0.001$, *n* = 3). **(B)** Mean P2/P1 ± SEM vs. IPI in control conditions (*black*) or with lidocaine present (*grey*).

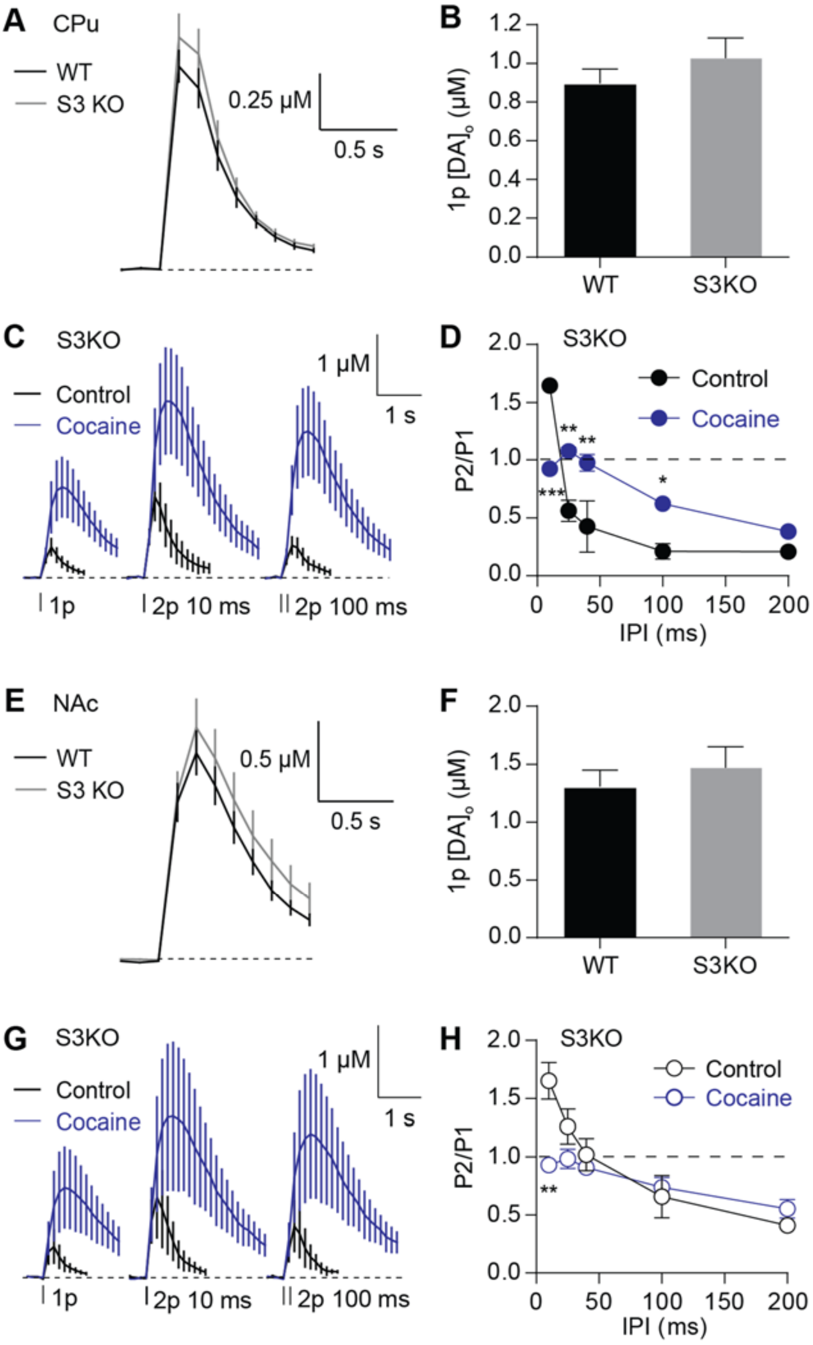


**Figure S5. The effects of cocaine on short-term plasticity are not dependent on synapsin III. (A, E)** Mean profiles of $[DA]_o$ transients (± SEM) evoked by single pulses in slices from wild-type (*black*) and S3KO (*grey*) animals in CPu (A, *n* = 6) and in NAc (E, *n* = 6). **(B, F)** Mean peak $[DA]_o$ (± SEM) following single pulses in wild-type (*black*) or S3KO (*grey*) animals in CPu (B) and in NAc (F). **(C, G)** Mean profiles of $[DA]_o$ transients (± SEM) evoked by single or paired pulses in control conditions (*black*) or cocaine (*blue*) in S3KO animals in CPu (C) or in NAc (G). **(D, H)** Mean P2/P1 (± SEM) vs. IPIs in control conditions (*black*) or in cocaine (*blue*) in CPu (D) and in NAc (H). Two-way ANOVA with Bonferroni's test for *post-hoc* comparisons; *$P < 0.05$, ** $P < 0.01$, *** $P < 0.001$.

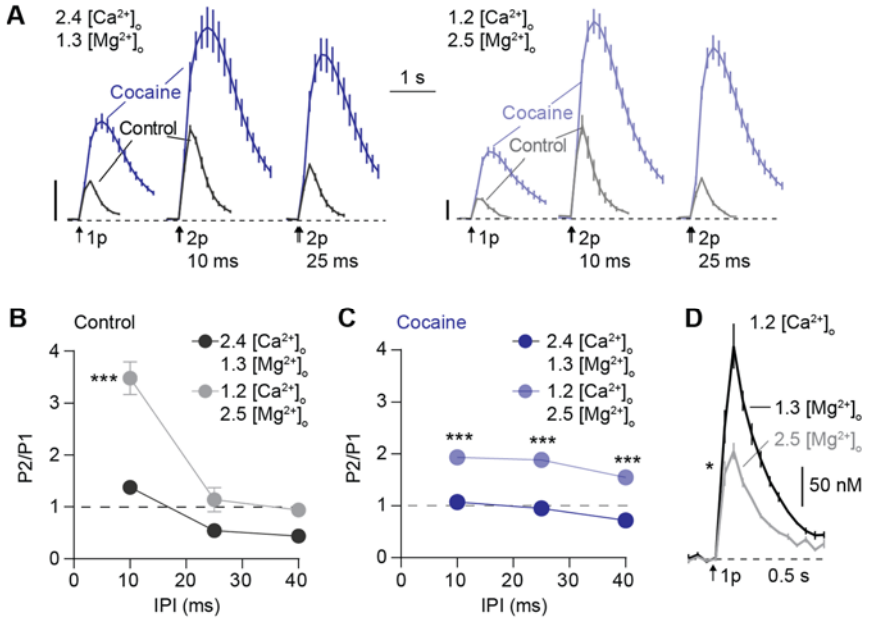


**Figure S6. $[Ca^{2+}]_o$ regulates short-term plasticity after DAT inhibition when divalent ion concentrations are maintained by $Mg^{2+}$; $Mg^{2+}$ also reduces DA release. (A)** Mean profiles of $[DA]_o$ transients (± SEM) evoked by 1 or 2 pulses in normal aCSF (2.4 mM $[Ca^{2+}]_o$/1.3 mM $[Mg^{2+}]_o$) (*left*) and in low $Ca^{2+}$ media (1.2 mM $[Ca^{2+}]_o$/2.5 mM $[Mg^{2+}]_o$) (*right*) in control conditions (*black*) or with 5 µM cocaine (*blue*). Data are normalised to 1p-evoked $[DA]_o$ (*vertical scale bars*) in normal aCSF. *n* = 4. **(B,C)** Mean P2/P1 (± SEM) vs. IPI after changes to $[Ca^{2+}]_o$ in control conditions (B) or cocaine (C). ****P*<0.001, *n = 4*. **(D)** Mean profiles of $[DA]_o$ transients (± SEM) evoked by single pulses in aCSF containing 1.2 mM $[Ca^{2+}]_o$ and either 1.3 mM (*black*) or 2.5 mM $[Mg^{2+}]_o$ (*grey*). Increasing $Mg^{2+}$ by 1.2 mM on a background of constant $[Ca^{2+}]_o$ reduces DA release. Unpaired t-test **P*<0.05, *n* = 3.

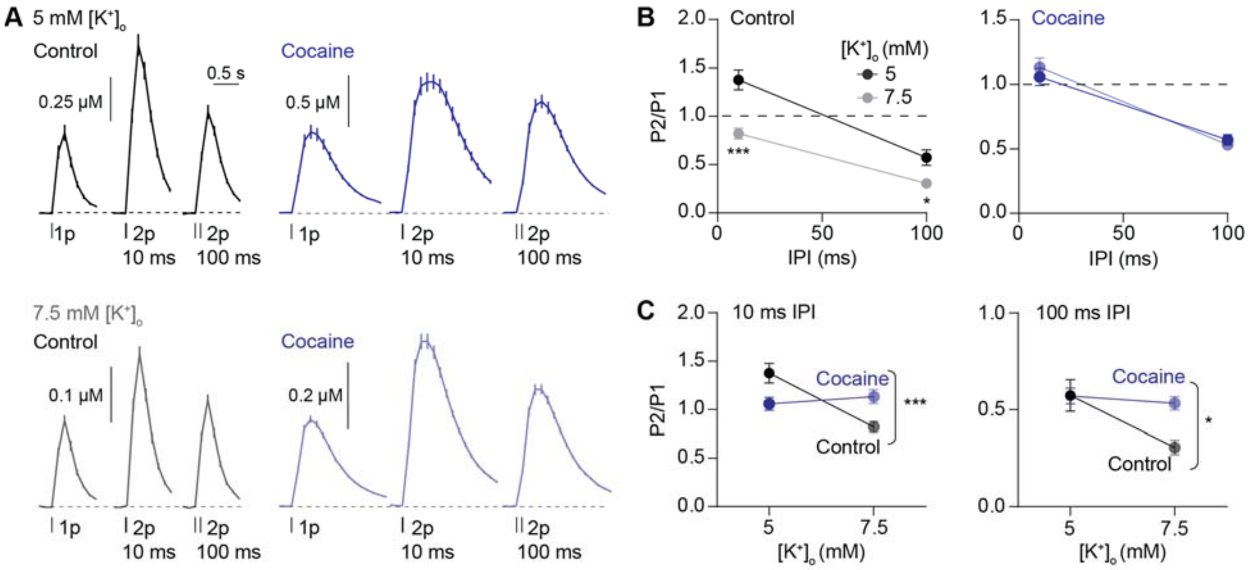


**Figure S7. Cocaine prevents enhancement of STD by increased $[K^+]_o$. (A**) Mean profiles (± SEM) of $[DA]_o$ transients in 5 mM (*upper*) and 7.5 mM $[K^+]_o$ (*lower*) in control conditions (*black-grey*) and in cocaine (*blue*). **(B)** Mean P2/P1 (± SEM) at IPIs of 10 ms and 100 ms in control conditions (*left*) and in cocaine (*right*) in 5 mM (*dark*) and 7.5 mM (*light*) $[K^+]_o$. **(C)** Mean P2/P1 (± SEM) in CPu at 10 ms IPI (*left*) and 100 ms IPI (*right*) in 5 mM and 7.5 mM $[K^+]_o$ in control conditions (*black-grey*) or with cocaine (*blue*). Two-way ANOVA with Bonferroni's test for *post-hoc* comparisons; * *P* < 0.05, *** *P* < 0.001.
